# Functional Characterization of Hsp110 in Drosophila Reveals its Essential and Dosage-Sensitive Role in Nervous System Integrity

**DOI:** 10.1101/2025.11.18.687301

**Authors:** Beatriz Rios, Shiyu Xu, Stephen M Farmer, Xin Ye, Lili Ye, Kevin A. Morano, Sheng Zhang

**Affiliations:** Center for Metabolic and Degenerative Diseases, The Brown Foundation Institute of Molecular Medicine, McGovern Medical School at the University of Texas Health Science Center at Houston (UTHealth Houston), Houston, Texas, USA; Program in Neuroscience, The University of Texas MD Anderson Cancer Center UTHealth Houston Graduate School of Biomedical Sciences (MD Anderson UTHealth Houston GSBS), Houston, Texas, USA; Program in Molecular and Translational Biology, The University of Texas MD Anderson Cancer Center UTHealth Houston Graduate School of Biomedical Sciences (MD Anderson UTHealth Houston GSBS), Houston, Texas, USA; Department of Microbiology & Molecular Genetics, McGovern Medical at the University of Texas Health Science Center at Houston (UTHealth Houston), Houston, Texas, USA; Program in Genetics and Epigenetics, The University of Texas MD Anderson Cancer Center UTHealth Houston Graduate School of Biomedical Sciences (MD Anderson UTHealth Houston GSBS), Houston, Texas, USA; Department of Neurobiology and Anatomy, McGovern Medical School at the University of Texas Health Science Center at Houston (UTHealth Houston), Houston, Texas, USA

## Abstract

The Hsp70 molecular chaperone system is the front line of defense in maintaining cellular proteostasis. In eukaryotes, ATP/ADP nucleotide exchange in the Hsp70 chaperone cycle is stimulated by Hsp110, a divergent member of the Hsp70 chaperone superfamily and co-chaperone of Hsp70. Hsp110 is also a known modifier of neurodegenerative and other protein misfolding-related disorders. Biochemical aspects of Hsp110 chaperone functions have been characterized *in vitro*, and pathway interactions have been extensively characterized genetically in yeast model systems; however, a detailed understanding of its physiological roles in metazoans, particularly in the nervous system has not been carried out. Taking advantage of the single Hsp110-encoding gene in the *Drosophila* genome, we conducted a comprehensive investigation of its expression and function in this animal model. Notably, *Drosophila* and human Hsp110 share significant similarity in their sequence, structure, and splicing variants. At the protein level, Hsp110 is ubiquitously expressed, with both cytosolic and nuclear distribution in a tissue-dependent manner. Functionally, while Hsp110 is dispensable for cell proliferation in developing larvae, it is essential for long-term cell survival and normal development of the nervous system, including non-autonomous effects on neuronal differentiation and glial cell migration. Furthermore, despite being identified as a potent suppressor of protein aggregation and neurotoxicity in multiple neurodegenerative diseases, higher levels of Hsp110 are detrimental in flies. Overexpression of Hsp40, another key co-chaperone of Hsp70, can mimic this effect. However, simultaneous overexpression of both Hsp40 and Hsp110 does not further exacerbate their detrimental effect. Together, these results demonstrate a critical role of Hsp110 in neuronal development and cell survival, and further suggest that *in vivo*, the levels and activities of Hsp110 and Hsp40 co-chaperones need to be properly balanced. Furthermore, this work supports the contention that the Hsp70 chaperone network must be considered as a whole when targeted for potential therapeutic purposes to meet the complex pathophysiological demands in multicellular organisms.

## Introduction

The cellular proteostasis network (PN) entails self-correction, clearance, and recycling mechanisms for proteins and is critical for many cellular functions, especially under stress conditions and for long-lived cells such as neurons. Growing evidence links dysfunction in the PN to diverse diseases from cancer to diabetes, and a large spectrum of brain degenerative diseases such as amyotrophic lateral sclerosis (ALS), Alzheimer’s (AD), Parkinson’s (PD), and Huntington’s (HD) diseases. These neurodegenerative diseases are characterized pathologically by the abnormal accumulation of specific disease-linked, misfolded proteins in neural tissue (Koszla and Solek 2024; Wilson et al. 2023). Thus, a clear understanding of the cellular PN is critical for developing efficient therapeutics against these debilitating diseases.

A cornerstone of the PN is the armamentarium of protein molecular chaperones that shepherd nascent polypeptides emerging from the ribosome, shield misfolded proteins from aggregation, and assist in targeting terminally damaged proteins for degradation. Key among these is the Hsp70 chaperone class, abundant and powerful protein remodeling engines present in multiple cellular compartments (Rosenzweig et al. 2019; Mayer and Bukau 2005). Powered by conformational changes resulting from ATP hydrolysis via its intrinsic ATPase activity, Hsp70 binds to nascent and misfolded proteins and facilitates their solubility and proper folding into functional conformations. In eukaryotic cells, a full Hsp70 chaperone cycle is facilitated by two key co-chaperones: DnaJ-domain containing Hsp40 that stimulates substrate binding to Hsp70 and its innate ATPase activity, and Hsp110 that acts as a nucleotide exchange factor (NEF) to facilitate ADP to ATP exchange in substrate-bound Hsp70, leading to subsequent substrate release and the resetting of the reiterative chaperone cycle (Rosenzweig et al. 2019).

Notably, Hsp110 itself is a member of the Hsp70 superfamily, sharing similar structure and domain architecture, including an ATP-binding N-terminal nucleotide binding domain (NBD) and a C-terminal substrate binding domain (SBD). The latter can be further divided into a β-sandwich subdomain (SBDβ) capable of polypeptide binding and an α-helical subdomain (SBDα) involved in regulating the binding and release of the peptide/protein substrate (Oh et al. 1999; Shaner and Morano 2007). Unlike classical Hsp70 chaperones, Hsp110 does not enhance protein refolding by itself despite harboring intrinsic ATPase activity (Andreasson et al. 2008; Raviol, Bukau, and Mayer 2006). Instead, it primarily acts as an NEF for Hsp70 (Dragovic et al. 2006; Raviol et al. 2006; Shaner, Sousa, and Morano 2006), as a “holdase” to passively bind unfolded or misfolded proteins thereby preventing their aggregation, and more prominently, as a key component of the potent Hsp110/Hsp70/Hsp40 disaggregase machinery that can dismantle existing aggregates to promote refolding or degradation (Rampelt et al. 2012; Shorter 2011; Wentink et al. 2020; Nillegoda and Bukau 2015).

In line with its key role in the chaperone cycle, in an earlier genome-wide RNAi screen for modifiers of aggregate formation by polyglutamine-expanded mutant Huntingtin (mHTT), the protein product of the HD gene, we identified Hsp110 together with Hsp40 and HSF1, the latter a key transcription factor that controls the stress-induced expression of Hsp70, as the most potent suppressors of mHTT aggregation. Additionally, altering Hsp110 expression levels was found to modulate neurodegeneration in an established fly HD model (Zhang et al. 2010). In addition to mHTT, Hsp110 has been shown to affect the folding and aggregation dynamics of a diverse range of misfolding-prone proteins, including Sup35 prion in budding yeast (Sadlish et al. 2008), other polyglutamine disease proteins, heat-denatured reporters, CF protein CFTR, amyloid beta (Aβ) in different model systems from *C elegans* to cultured mammalian cells (Kuo et al. 2013; Aguado et al. 2015; Yamagishi et al. 2010; Ishihara et al. 2003; Rampelt et al. 2012; Montresor et al. 2025; Saxena et al. 2012), mutant SOD1 in mouse and squid axoplasm-derived ALS models (Wang et al. 2009; Song et al. 2013), α-Synuclein amyloid fibrils *in vitro,* and in mouse PD models (Gao et al. 2015; Taguchi et al. 2019). Most notably, in recent genetic and RNAseq studies on human AD samples, Hsp110 together with Hsp40 and Hsp70 stand out as AD-resilience genes, showing consistent upregulation in neurons from AD-resilient brains and downregulation in AD-affected brains (Llibre-Guerra et al. 2025; Castanho et al. 2025). Together, these reports support a critical role for Hsp110 and the Hsp70 chaperone machinery in protein misfolding diseases across evolution.

Hsp110 is conserved in all eukaryotes. However, noticeable structural and functional differences exist between Hsp110 orthologs in single-cell species and those of multicellular organisms. Specifically, when compared to the protein products of the two yeast Hsp110 genes, *SSE1* and *SSE2,* metazoan Hsp110 harbors two significantly longer extensions: a loop insertion within the SBDβ subdomain and an elongated C-terminal tail following the SBDα subdomain. Furthermore, a recent analysis identified a novel intrinsically disordered region (IDR) conserved at the extreme C-terminus of both fly and human Hsp110, but absent in yeast counterparts (Yakubu and Morano 2021). Importantly, in *in vitro* assays, both the fly and human IDRs were sufficient to prevent the aggregation of Aβ peptides (Yakubu and Morano 2021). Together, these observed differences point to a potential functional divergence with different regulatory mechanisms of metazoan Hsp110s from those in single-cell species to accommodate the more complex physiology of multicellular organisms.

Although metazoan Hsp110s have been extensively characterized biochemically, there are limited *in vivo* studies on their physiological roles, partially due to the presence of three Hsp110 genes (Hsp105 (HSPH1), APG-1 (HSPA4L), APG-2 (HSPA4)) with overlapping expression patterns in mammals that complicate their functional study (Easton, Kaneko, and Subjeck 2000; Shaner and Morano 2007). In mice, APG-1 and APG-2 double-knockouts (KO) led to smaller lungs, respiratory stress, and neonatal death (Mohamed et al. 2014), while mice with Hsp105 single KO were viable but showed increased levels of Tau hyperphosphorylation and amyloid accumulation, the hallmarks of AD brains (Eroglu, Moskophidis, and Mivechi 2010). Despite the extensive biochemical characterization of Hsp110 and analysis of biological roles in budding yeast, and the clear implications for Hsp110’s importance in defense against protein misfolding consequences in neurons, Hsp110’s effect on animal development at the cellular level, especially during neuronal differentiation, remains largely unexplored.

*Drosophila* has a single *hsp110* gene (also known as *hsc70Cb, dhsp110,* and *hsc70,* which we will refer to as *hsp110* for the gene and Hsp110 for its protein products hereafter), sharing ∼40% identity and ∼60% similarity to vertebrate Hsp110 at the amino acid level (Easton, Kaneko, and Subjeck 2000; Zhang et al. 2010). We therefore exploited this feature in the genetically tractable fly model to carry out a detailed characterization of Hsp110 with an emphasis on its roles in the nervous system. Our results revealed remarkable conservation of the fly Hsp110 with its human counterparts at multiple levels, from gene structure to conformation and splicing variants, as well as its essential and dosage-sensitive effects on cell survival and nervous system integrity. In particular, Hsp110 demonstrated to be dispensable for cell proliferation but critical for long-term cell survival and proper development of the nervous system. However, its overexpression was detrimental, despite extensive evidence supporting its essential role in preserving the PN. These findings suggest the existence of robust regulatory mechanisms, potentially mediated through the metazoan-specific divergent domains in Hsp110s, to efficiently modulate its endogenous activity for productive Hsp70 chaperone cycles in response to complex physiological and pathological demands *in vivo*.

## Results

### Structural and sequence similarity of *Drosophila* and mammalian Hsp110 proteins

The FlyBase documented eight transcript isoforms from the *Drosophila hsp110* gene, which encodes shorter (804; designated as Hsp110-A in the Flybase) or longer (836; Hsp110-G) amino acid-long protein products differing by a differentially spliced small coding exon 5 encoding a 32 amino acid-long insertion (Fig. 1A) (http://flybase.org/reports/FBgn0026418#expression). Interestingly, similar splicing isoforms (HSP105A and HSP105B) also exist in humans and mouse HSP105 that differ by a ∼44 amino acid-long insertion (Fig. S1). Moreover, an AlphaFold 3 prediction revealed that these extra sequences, all derived from alternatively spliced exons, similarly contribute to an unstructured, elongated region within the loop insertion in the SBDβ subdomain (arrows in Fig. 1B).

**Fig 1.**
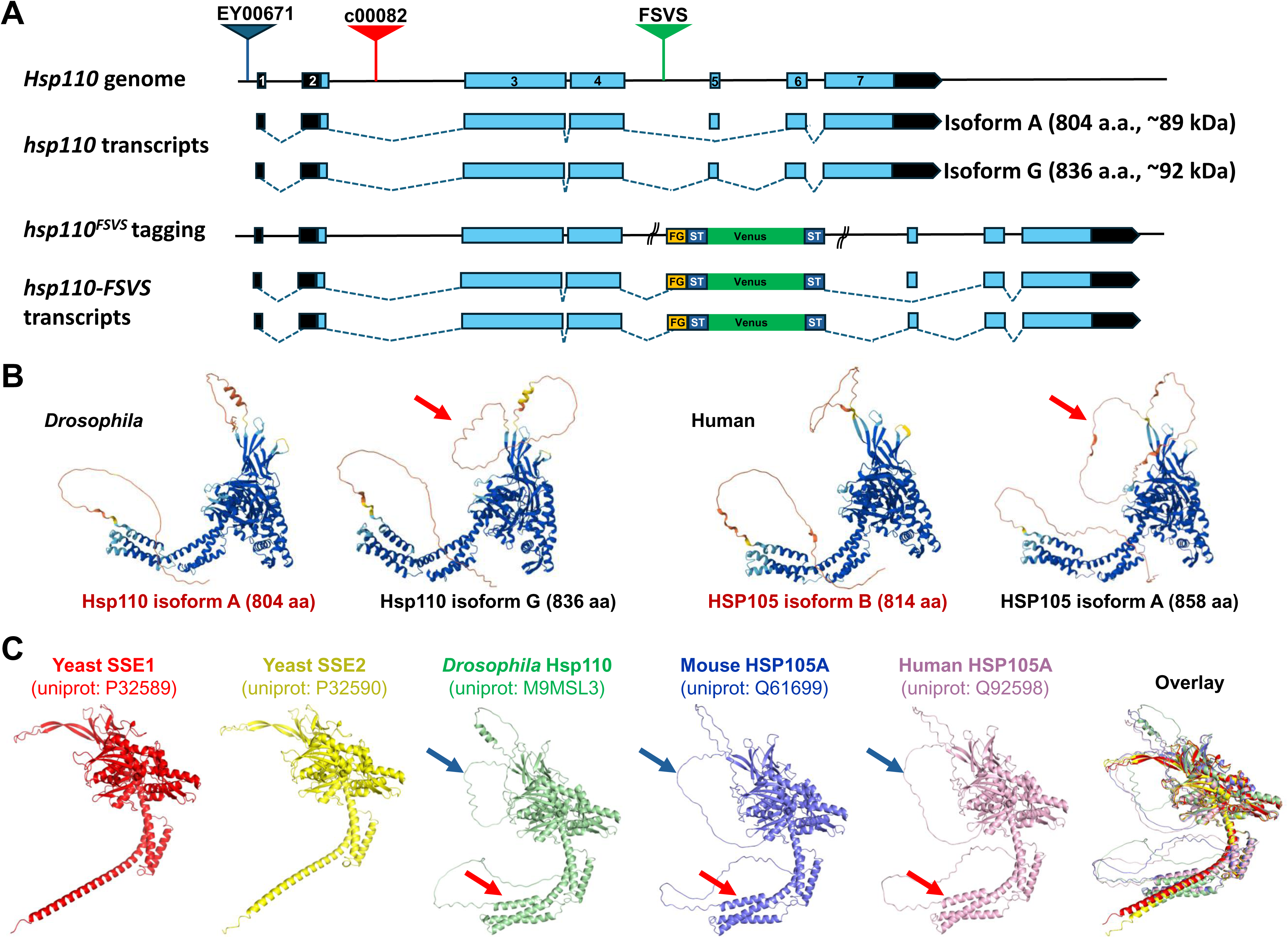
*Drosophila hsp110* genome and the structural comparison of Hsp110 proteins across species. (A) Cartoon illustration of the *hsp110* genomic structure, associated insertional alleles (colored triangles), the A and G transcription isoforms encoding the short and long Hsp110 protein products, respectively, and the predicted genomic structure and transcript isoforms from the *hsp110^FSVS^* gene trap lines, all drawn in scale except for the *hsp110^FSVS^* lines, for which only the artificial exon encoding the 3x-FLAG-Venus-StrepTagII (FSVS) tags was shown, with the curved parallel lines representing the surrounding sequences from the host piggyBac transgenic vector. FG: 3xFLAG tag. ST: StrepTagII. Lines depict introns, black boxes for the non-coding regions, and blue boxes for the coding regions. (B) Structural comparison of fly Hsp110 A and G isoforms with human HSP105 Alpha (HSP105A) and Beta (HSP105B) isoforms, modeled by AlphaFold. Red arrows highlight the extended loop insertion from the alternatively spliced exons in the SBDβ subdomain. (C) Structural comparison of Hsp110 proteins from different species, including SSE1 and SSE2 from yeast *S cerevisiae*, Hsp110 in *Drosophila*, HSP105-alpha (Hsp105A, HSPH1) from mouse and human, and their overlay, as predicted by AlphaFold 3. Blue arrows highlight the unstructured loop insertion between the SBD, red arrows highlight an additional Helix-Turn-Helix motif and the long, unstructured C-terminus tail specifically in metazoan Hsp110 proteins.

At both sequence and structural levels, *Drosophila* Hsp110 shows more similarities with mammalian Hsp110s than with yeast Hsp110s (*SSE1* and *SSE2* from *S. cerevisiae*, and *PSS1* in *S. pombe*). For example, the linker sequence that connects the NBD and SBD domains is the highly distinctive PFKFED in yeast Hsp110s (Chakafana et al. 2021), EFGVTD in fly, and EFSV(I)TD in human Hsp110s (Fig. S1). Moreover, at the structural level, the AlphaFold 3 prediction revealed several unique features in fly and mammalian Hsp110s that distinguish them from yeast Hsp110s: (1) the distinctively long unstructured loop insertion within the β-sandwich SBDβ (blue arrows in Fig 1C. ∼45 amino acids in fly Hsp110-A isoform and ∼77 amino acids in the Hsp110-G isoform; ∼45 amino acids in human Hsp105B and ∼89 amino acids in human Hsp105A); (2) an additional helix-turn-helix motif that is adjacent to and mirrors the SBDα lid subdomain (red arrows in Fig 1C); (3) an extended and unstructured C-terminal IDR tail (Fig. 1C and Fig. S1). At the protein sequence level, among the three human Hsp110 proteins, *Drosophila* Hsp110-G isoform is most similar at the amino acid level to human APG1 (HSPA4L, 43% identity, 62% Similarity) and APG2 (HSPA4, 43% Identity, 60% Similarity), followed by HSP105A (HSPH1, 40% Identity, 59% Similarity), as analyzed with DIOPT (https://www.flyrnai.org/cgi-bin/DRSC_orthologs.pl).

### Characterization of Hsp110 mutant alleles and genome-tagging lines

To study the physiological role of Hsp110 in *Drosophila*, we characterized several transposon alleles inserted in the *hsp110* genome region (Fig. 1A). Complementation testing revealed that two alleles, *c00082* and *l(3)s64906* formed a complementation group. They were homozygous lethal but viable when in trans-heterozygous with the other insertional lines (summarized in Table 1). In addition, two gene trap alleles, KSC-115570 and KSC-115275 (“FSVS” in Fig. 1A), were similarly inserted inside the intron between the coding exon 4 and the alternatively spliced exon 5. These two gene trap lines carried an artificial exon encoding a tripartite tag, an in-frame fusion of three copies of the FLAG epitope, and a fluorescent Venus protein flanked by two copies of the StrepTagII tag (3xFLAG-StrepTagII-Venus-StrepTagII, abbreviated as “FSVS” hereafter), flanked by splicing acceptor and donor sequences (Fig. 1A) (Lowe et al. 2014). Accordingly, if the artificial exon was transcribed, spliced, and translated as part of the native *hsp110* gene, then the FSVS tripartite tag became part of the extended unstructured loop insertion in the SBDβ subdomain, producing endogenously tagged Hsp110-FSVS fusion protein products, with their expression under the control of endogenous regulatory elements of the native *hsp110* gene. We observed that the original KSC-115570 and KSC-115275 lines were both homozygous lethal. After backcrossing the KSC-115570 line with the wildtype (WT) line for several generations, it became homozygous viable, suggesting that the original lethality was due to background mutations in the parental line that were removed in subsequent rounds of crosses. For simplicity, the KSC-115570 and KSC-115275 lines are abbreviated as *hsp110^FSVS1^* and *hsp110^FSVS2^*, respectively, hereafter.

**Table-1.**
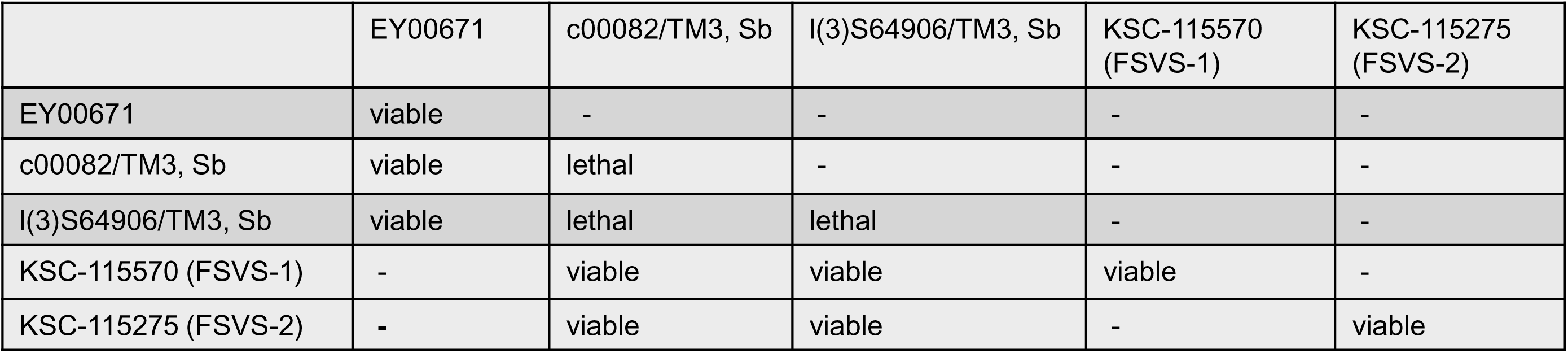
Summary of complementation tests of Hsp110 alleles.

|  | EY00671 | c00082/TM3, Sb | l(3)S64906/TM3, Sb | KSC-115570 (FSVS-1) | KSC-115275 (FSVS-2) |
| --- | --- | --- | --- | --- | --- |
| EY00671 | viable | - | - | - | - |
| c00082/TM3, Sb | viable | lethal | - | - | - |
| l(3)S64906/TM3, Sb | viable | lethal | lethal | - | - |
| KSC-115570 (FSVS-1) | - | viable | viable | viable | - |
| KSC-115275 (FSVS-2) | - | viable | viable | - | viable |

To further characterize the above alleles and examine expression of the endogenous Hsp110 protein products, we generated specific rat anti-Hsp110 polyclonal antiserum. In immunoblot assays on homogenates from whole WT adult flies (lane 1 in Fig. 2A), the anti-Hsp110 antibody recognized a strong ∼95 kDa band and a slightly larger ∼100 kDa band; the latter was present at levels about one-sixth that of the ∼95 kDa product (quantification in Fig. 2B). These ∼95 and 100 kDa products likely represent the two predicted splicing isoforms of Hsp110 with expected sizes of ∼88kDa and 93kDa, respectively. In support of this prediction, in flies transgenic for the short A-isoform of Hsp110, when the transgene expression was induced by a mild *armadillo*-GAL4 (*arm*-GAL4) (lanes 2 and 4 in Fig. 2C) or by a stronger *daughterless*-GAL4 (*da*-GAL4) (lane 2 in Fig. 2G) drivers, only the level of the ∼95 kDa product became proportionally higher, confirming that the ∼95kDa band was the product of the short Hsp110 A isoform. Importantly, on homogenates from adult flies homozygous for the *hsp110^FSVS1^* gene trap line, both the ∼95 kDa and ∼100 kDa bands were absent and replaced by a stronger ∼130 kDa band and a slightly larger but significantly weaker band just above it (see the magnified view of the band area highlighted by dashed line in Fig. 2A), mirroring the pattern of the two endogenous Hsp110 protein isoforms. Moreover, a similar 130 kDa band with comparable expression level was also present in *hsp110^FSVS2^* flies as in *hsp110^FSVS1^* (Fig. 2D). Lastly, the ∼130 kDa molecular masses were consistent with the predicted sizes of the Hsp110-FSVS fusion and could be simultaneously detected by anti-Hsp110 with anti-FLAG (Fig. 2A) and anti-GFP (Fig. 2D) antibodies. Together, these results support that in the *hsp110^FSVS1^* and *hsp110^FSVS2^* gene trap lines, the endogenous ∼95kDa and ∼100 kDa Hsp110 protein products were replaced by two larger ∼130 kDa fusions that contain the in-frame FSVS tripartite tags. Further quantification revealed that in homozygous *hsp110^FSVS1^* adults, the total level of ∼130 kDa Hsp110-FSVS fusions was about 70% that of endogenous ∼95 and 100 kDa Hsp110 proteins in WT flies (Fig. 2A, 2D, and quantification in Fig. 2E), indicating a mild diminishment of Hsp110 expression levels by the inserted artificial exon. Nevertheless, as the backcrossed *hsp110^FSVS1^* gene trap line was homozygous viable and fertile, these results support the conclusion that the inclusion of the FSVS tag in the unstructured loop did not disrupt normal function of Hsp110, and the resulting Hsp110-FSVS fusions were fully functional. As their expression was under the control of native *hsp110* regulatory elements, Hsp110-FSVS fusions were likely expressed with patterns mirroring those of the endogenous Hsp110. The *hsp110^FSVS^* gene trap lines thus provided ideal tools for detecting and manipulating the endogenous Hsp110 proteins.

**Fig 2.**
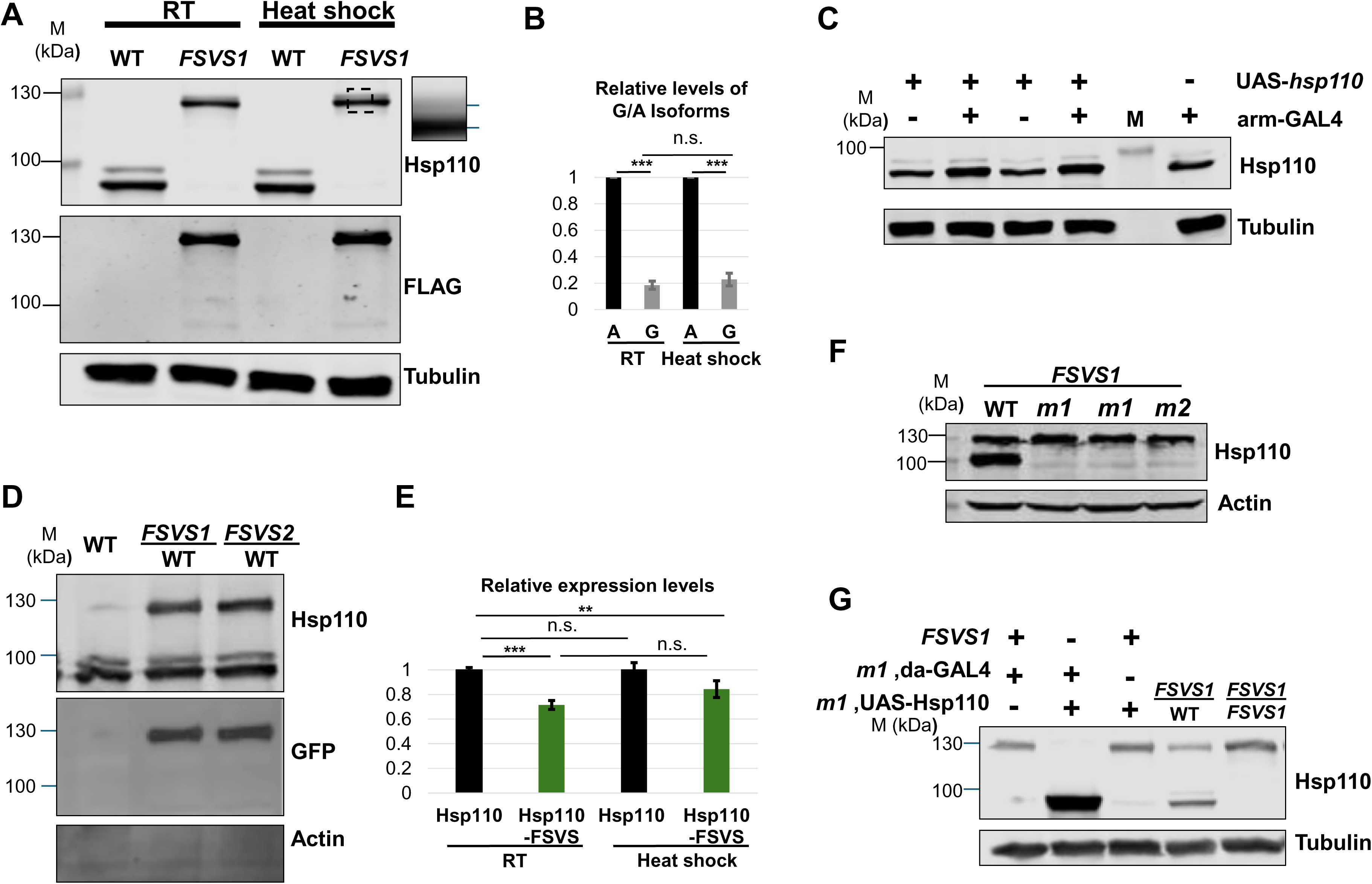
Hsp110 protein products in WT and *hsp110* insertional alleles. Western blot analyses for Hsp110 proteins in different fly lines, with Actin or Tubulin as loading controls, as annotated. (A) Two Hsp110 protein isoforms exist in *Drosophila*. Immunoblot on homogenates of adult flies from WT (genotype: *w^1118^*) or homozygous *hsp110^FSVS1^* endogenous tagging line (FSVS1. Genotypes: *w^1118^*; *hsp110^FSVS1^*/*hsp110^FSVS1^*), cultured at room temperature (RT) or after one-hour heat-shock at 37°C water bath followed by one hour of recovery at room temperature (Heat shock), sequentially probed with anti-Hsp110 (top) and anti-FLAG (middle panel) antibodies. The small right panel is a zoomed-in view of the boxed area of the ∼130kDa Hsp110-FSVS bands in lane 4 from *hsp110^FSVS1^* flies, revealing a slightly larger but weaker band on top of the dominant ∼130 kDa band, mirroring the pattern of that in WT samples. (B) Bar graph presentation of the relative protein levels of Hsp110 A and G isoforms in WT adult flies, cultured at room temperature (RT) or after one-hour heat-shock (Heat shock), normalized against loading control Tubulin. *** p<0.001 (n= 5 independent repeats). (C) The major ∼95 kDa product corresponds to Hsp110 A isoform. Immunoblot for Hsp110 proteins in adult flies carrying the transgenes for UAS-*hsp110*-A isoform alone (genotype: UAS-*hsp110*/+), *arm*-GAL4 driver alone (genotype: *arm*-GAL4/+), or both (genotype: *arm*-GAL4/+; UAS-*hsp110*/+), as annotated. The levels of ∼95 kDa product, but not the larger ∼100 kDa band, showed a mild increase only in flies carrying both UAS-*hsp110* and *arm*-GAL4 driver (lanes 2 and 4). (D) Both *hsp110^FSVS1^* and *hsp110^FSVS2^* lines express the same endogenously tagged Hsp110-FSVS fusion protein products. Immunoblot for Hsp110 (top panel) and GFP (middle panel) on homogenates from adults of WT control (genotype: *w^1118^*) or flies heterozygous for *hsp110^FSVS1^* (FSVS1. Genotypes: *w^1118^*; *hsp110^FSVS1^*/*+*) or *hsp110^FSVS2^* (FSVS2. Genotypes: *w^1118^*; *hsp110^FSVS2^*/*+*) tagging lines over the WT chromosome. (E) Bar graph presentation of the relative total levels of ∼95 kDa Hsp110 and ∼130 kDa Hsp110-FSVS protein products in WT and homozygous *hsp110^FSVS1^*adult flies, respectively, cultured at room temperature (RT) or after one-hour heat-shock, normalized against loading control Tubulin. ** p<0.01 *** p<0.001 (n= 6 independent repeats). (F) Absence of endogenous Hsp110 proteins in *hsp110^m1^* and *hsp110^m2^* alleles. Immunoblot for Hsp110 in homogenates from adult flies heterozygous for *hsp110^FSVS1^* tagging line over WT control (lane 1), *hsp110^m1^* (lanes 2 and 3 as two independent repeats), or *hsp110^m2^* (lane 4) alleles, as annotated. (G) The lethality of homozygous *hsp110^m1^* flies is due to the loss of *hsp110* protein products. Immunoblot for Hsp110 in homogenates from adult flies of the annotated genotypes. (lane 1) Control of the heterozygous *hsp110^FSVS1^* tagging line over the recombinant line of *hsp110^m1^* allele with *da*-GAL4 driver (genotypes: *w^1118^*; *hsp110^FSVS1^*/ *da*-GAL4, *hsp110^m1^*). (lane 2) Homozygous *hsp110^m1^* adults rescued by the ectopically expressed Hsp110-A isoform from UAS-*hsp110* transgene driven by *da*-GAL4 (genotype: “*w^1118^*; *da*-GAL4, *hsp110^m1^*/UAS-*hsp110*, *hsp110^m1^*”). (lane 3) Control of heterozygous *hsp110^FSVS1^*tagging line over the recombinant line of *hsp110^m1^* allele with UAS-*hsp110* transgene (genotype: “*w^1118^*; *hsp110^m1^*, UAS-*hsp110* /*hsp110^FSVS1^*”). (lane 4) Heterozygous *hsp110^FSVS1^* over WT (genotypes: *w^1118^*; *hsp110^FSVS1^*/+). (lane 5) Homozygous *hsp110^FSVS1^* tagging line. Notice the absence of endogenous ∼95/100 kDa Hsp110 products in lanes 1, 3, and 5, and the significantly higher levels of ectopically expressed ∼95kDa Hsp110 product in homozygous *hsp110^m1^* adults (lane 2).

Lastly, we tested whether the expression of endogenous Hsp110 was inducible in response to heat shock stress by incubating WT and homozygous *hsp110^FSVS1^* adult flies in a 37°C water bath for one hour, followed by recovery at room temperature for another hour. Immunoblotting on homogenates from these flies did not reveal a significant increase in the total levels of endogenous Hsp110 or Hsp110-FSVS fusion proteins (Fig. 2A and quantification in Fig. 2E), suggesting that Hsp110 was not a robust, heat stress-inducible gene under the tested conditions.

### Hsp110 is essential for Drosophila development

*c00082* and *l(3)s64906* lines proved to be in the same complementation group (Table 1). Both were embryonic lethal as homozygotes but fully viable when in trans-heterozygous with either of the two *hsp110^FSVS^* gene trap lines (e.g., genotype “*c00082/hsp110^FSVS1^*”). In immunoblot assays on homogenates from these trans-heterozygous adults, only the ∼130 kDa Hsp110-FSVS fusion proteins were present, while the endogenous ∼95 kDa and ∼100 kDa bands were mostly absent (Fig. 2F), supporting that both *c00082* and *l(3)s64906* insertions caused a significant loss of the endogenous protein products of the *hsp110* gene. Importantly, the lethality of homozygous *c00082* flies could be rescued by the ectopically expressed Hsp110-A isoform from UAST-based transgenes directed by a ubiquitous *da*-GAL4 driver (lane 2 in Fig. 2G). The rescued homozygous *c00082* flies showed no apparent morphological defects as adults, thus confirming that the embryonic lethality was due to the loss of endogenous Hsp110. We observed that the rescued adult females were sterile. Further dissection of these flies revealed severely abnormal ovarian development, characterized by ovaries that were largely devoid of mature egg chambers. In contrast, the ovaries of age-matched WT controls were significantly larger and contained egg chambers at various maturation stages (Fig. S2). The sterile defect is likely due to a lack of UAST-based transgenes expression in the female germline (Rorth 1998). Together, the results demonstrate that Hsp110 is critical for *Drosophila* development and ovary growth, with *c00082* and *l(3)s64906* lines as genetically, useful strong loss-of-function alleles, abbreviated as *hsp110^m1^* and *hsp110^m2^*, respectively, hereafter.

### Ubiquitous expression and subcellular localizations of Hsp110 proteins

We validated that the rat anti-Hsp110 antibodies we generated could detect the endogenous Hsp110 protein in immunofluorescence staining. Specifically, in FRT-mediated mosaic clones composed of cells homozygous for either of the two *hsp110* mutant alleles (see the section below), the signal intensity from anti-Hsp110 antibody staining was absent when compared to the surrounding heterozygous cells (Fig. S3A, B, with more details below). Moreover, in tissues from the *hsp110^FSVS^* gene trap lines, simultaneous staining with anti-Hsp110 and anti-GFP antibodies revealed identical expression and subcellular patterns of the Hsp110-FSVS fusion (Fig. 3A-D).

**Fig 3.**
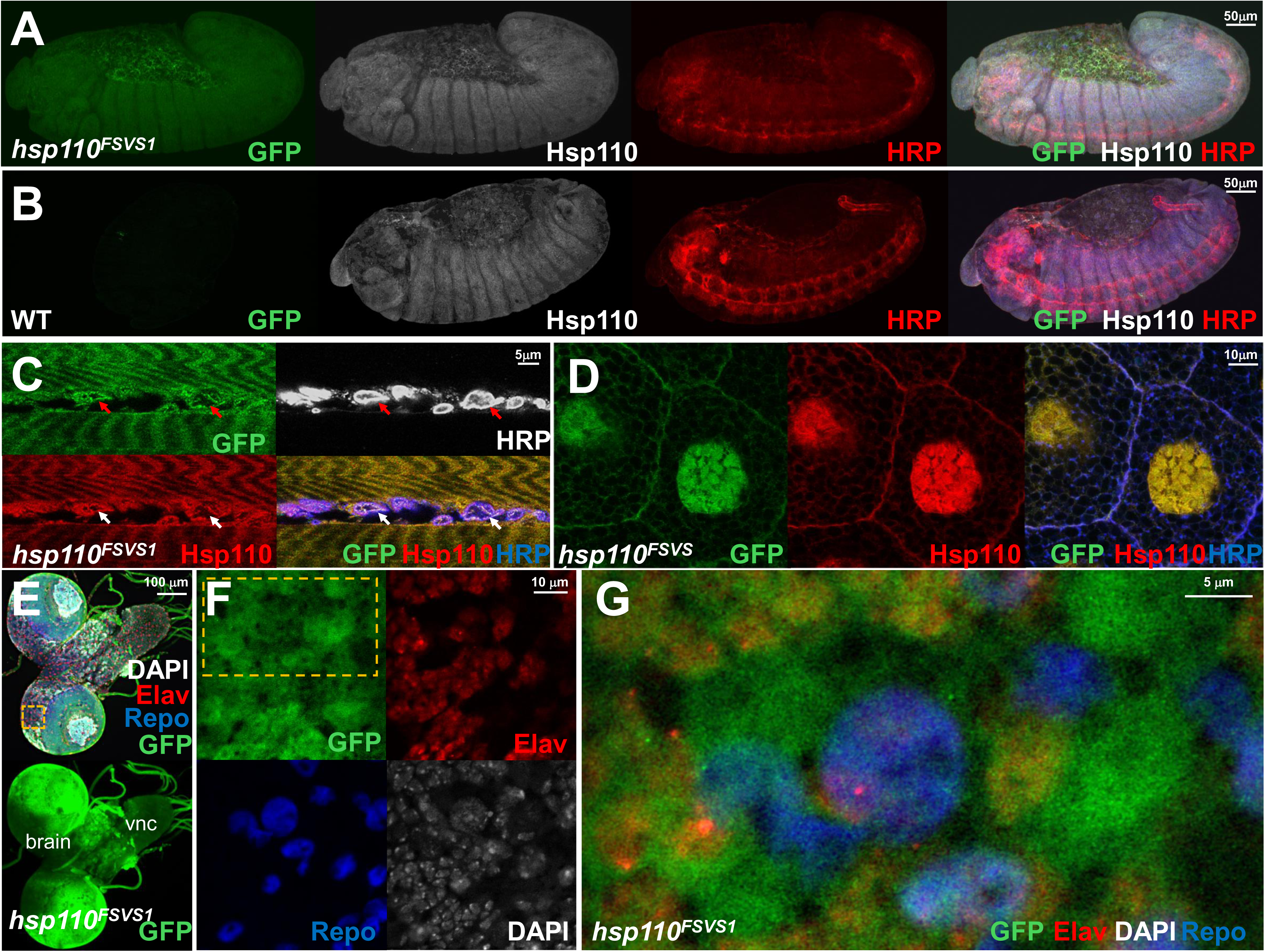
Ubiquitous expression and discrete subcellular localization of Hsp110 proteins in Drosophila. Confocal imaging for the expression and subcellular localization of Hsp110 together with cellular markers in different tissues and developmental stages, shown in single-channel or overlaying images, as annotated. (A, C, D) *hsp110^FSVS1^*tagging or (B) WT flies, co-labeled with antibodies against Hsp110, GFP for the Venus tag in the Hsp110-FSVS fusion, and HRP for neuronal (A-C) and cellular (D) membrane, as annotated. (A, B) Stage 12 embryos from (A) *hsp110^FSVS1^*and (B) WT flies. (C) Muscle and neuromuscular junctions (NMJs, highlighted by arrows) and (D) salivary gland cells from third instar larvae. (E-G) Brain and ventral nerve cord (vnc) from third instar larvae, triple-labeled for antibodies against GFP to detect the Venus tag in the Hsp110-FSVS (green), Elav (marker for neuronal nucleus, red), Repo (marker for glial nucleus, blue), and DAPI for DNA (white), as annotated. (F, G) High magnification view of the boxed area in the protocerebral region in (E) or (F), respectively, in the larval brain from the *hsp110^FSVS1^* genome-tagging line. Note that the Hsp110-FSVS fusion is present in all the cells, predominantly as puncta in the cytoplasm.

Using the validated anti-Hsp110 antibodies and the Hsp110-FSVS genome tagging lines, we next examined the distribution of Hsp110 protein during development and in different tissues, which revealed a ubiquitous and predominantly cytoplasmic presence of Hsp110. For example, in stage 12 embryos from *hsp110^FSVS1^*gene trap lines, staining with anti-GFP and anti-Hsp110 antibodies showed an identical, ubiquitous presence of Hsp110 in all the examined cells, with no apparent enrichment in particular tissues (Fig. 3A). An almost identical ubiquitous expression pattern was also observed in WT stage 12 embryos stained by anti-Hsp110 (Fig. 3B). Similarly, in larval and adult males and females, Hsp110 was ubiquitously expressed in all examined tissues, including neurons and glial cells in the brain (Fig 3C-G and data not shown).

At subcellular levels, Hsp110 predominantly localized in the cytoplasm, however we observed variable levels of nuclear distribution in different cell types. For example, in muscles of third instar larvae, Hsp110 was enriched in the cytoplasm of muscle myofibers and neuromuscular junctions (NMJs) (arrows in Fig. 3C). In neurons and glial cells of third instar larval brain, Hsp110 predominantly localized to the cytoplasm, but could also be detected in the nucleus, as revealed by its partial colocalization with Elav and Repo, two markers for the nucleus of neurons and glial cells (Alfonso and Jones 2002; O’Neill et al. 1994), respectively (Fig. 3E, F). However, in polyploid salivary gland cells, in addition to its cytoplasmic localization, Hsp110 also associated with the plasma membrane and, more strikingly, showed strong enrichment in the nucleus (Fig. 3D). Closer inspection revealed that at subcellular levels, Hsp110 was not evenly dispersed inside cells, but nucleated in tiny puncta (Fig. 3G).

### Hsp110 is essential for tissue development and long-term cell survival

Hsp110’s physiological roles in animal development, such as cell proliferation, differentiation, and survival, especially in neurons, are poorly understood. Because it was not possible to address these questions in homozygous *hsp110* mutants, which are embryonic lethal, we used the Flippase (FLP)/ FLP Recognition Target (FRT) -inducible mosaic system to generate clones of cells homozygous for the mutants in an otherwise WT heterozygous background, thereby bypassing animal lethality (Chou and Perrimon 1992; Xu and Rubin 1993). We first utilized the heat shock-inducible MARCM system to generate mosaic clones, an approach that expresses the FLP ubiquitously in all tissues in a heat shock-dependent manner (“*hs-*FLP*”*), with homozygous mutant cells positively labeled by fluorescent GFP (Lee and Luo 1999). For both *hsp110^m1^* and *hsp110^m2^* alleles, the mosaic was induced by heat shock during the early second instar larval and then examined about 60 hours later. By late third instar, GFP-positive clones composed of large numbers of homozygous mutant cells were present in various tissues, including the brain, eye, and wing imaginal discs (Fig. S3 and 4A-C), suggesting that mutant cells could survive and proliferate during development. However, few of the surviving progeny were non-balancer flies (e.g., genotype “*hs-*FLP/+, tub>GFP; tub-Gal80, FRT80B/*hsp110^m2^*, FRT80B”), that generated mutant clones in various tissues during development. Almost all the surviving adults carried the balancer chromosome (e.g., genotype as “*hs-*FLP/+, tub>GFP; tub-Gal80, FRT80B/TM6B, Tb”), which did not harbor the FRT target site and therefore could not generate the FLP-induced mutant clones. Moreover, the few non-balancer escapers often had deformed adult structures, such as abnormally shaped eyes (Fig. 4F, H). Since the mosaic clones for mutant *hsp110* were induced by a *heatshock* promoter, that targeted all fly tissues, the low survival rate of non-balancer flies suggests Hsp110 is critical for the development and function of essential tissues. The widespread presence of mutant *hsp110* clones during development proved to be detrimental and more often than not, lethal.

**Fig 4.**
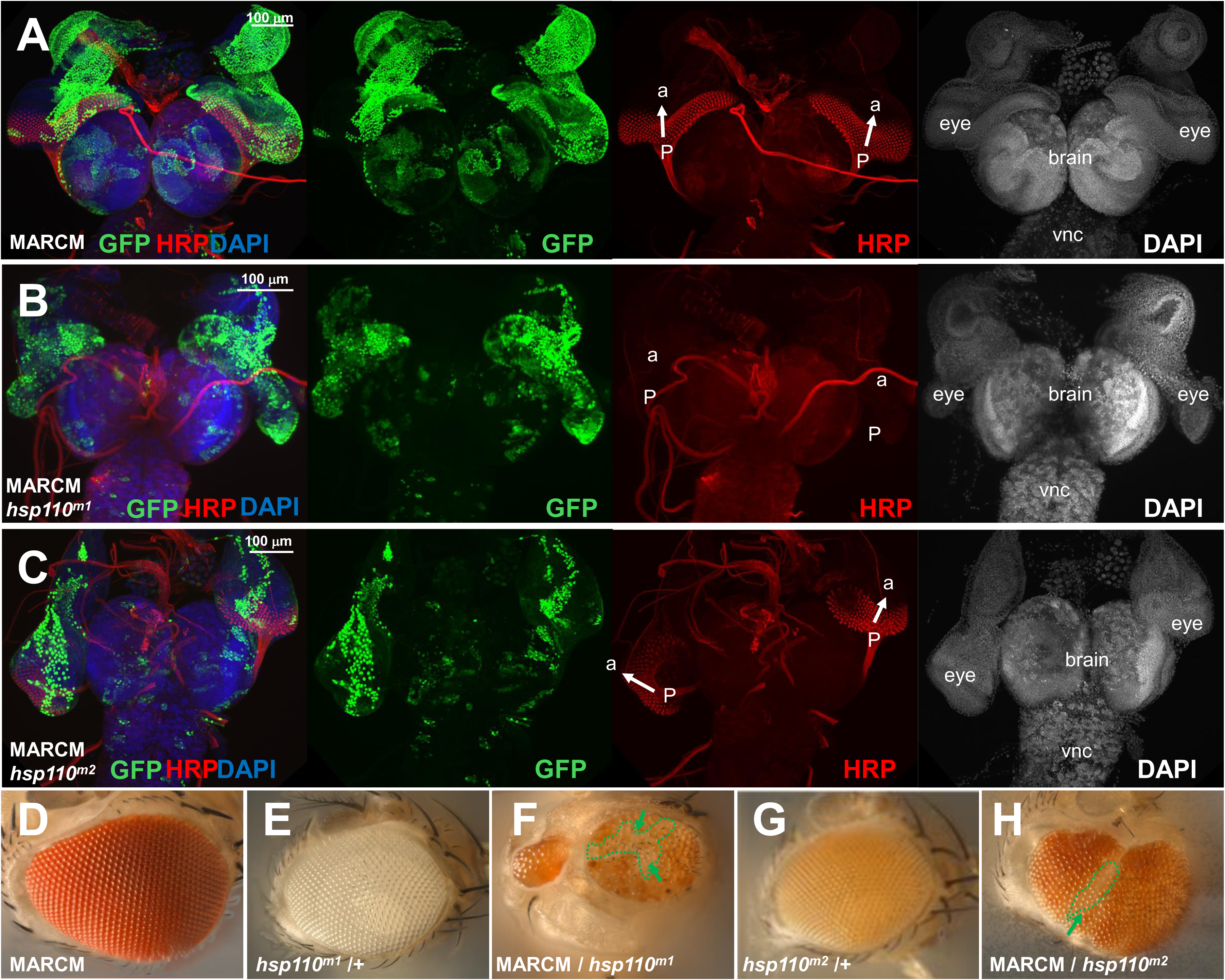
Hsp110 is dispensable for cell proliferation but essential for neuronal differentiation and cell survival. (A-C) Confocal images of brain and eye-antennal imaginal discs from third-instar larvae containing GFP-labeled MARCM clones (green) homozygous for (A) WT control, (B) *hsp110^m1^* or (C) *hsp110^m2^* mutants, co-stained for neuronal membrane marker HRP (red) and DAPI (blue in overlaying images and gray in single channel), as annotated. The differentiation of neuronal photoreceptors initiates from the posterior (p) of the eye imaginal disc, where it connects to the brain lobes through the optic stalk, and progressively moves to the anterior (a), as highlighted by white arrows. Notice the presence of large numbers of GFP-labeled homozygous mutant cells for both (B) *hsp110^m1^* and (C) *hsp110^m2^* alleles in both the eye-antennal discs as well as the brain and ventral nerve cord (vnc), and the presence of HRP-positive photoreceptor cells in the eye imaginal disc in (C) similar to that in WT (A), but absent in (B). (D-H) Bright-field images of adult eyes. (D) Parental WT MARCM line (genotype: “*hs*-FLP; MARCM, FRT80B/ MARCM, FRT80B”), which is bright red. (E, G) Parental lines of heterozygous (E) *hsp110^m1^* or (G) *hsp110^m2^*mutant over WT, which show white (E, genotype: “*w^1118^*; *hsp110^m1^*, FRT80B/+”) or orange (G, genotype: “*w^1118^*; *hsp110^m2^*, FRT80B/+”) color, respectively. (F, H) Flies with heat shock-induced MARCM mosaic clones for (F) *hsp110^m1^* (genotype: “*hs*-FLP; MARCM, FRT80B/ *hsp110^m1^*, FRT80B”) or (H) *hsp110^m2^*(genotype: “*hs*-FLP; MARCM, FRT80B/ *hsp110^m2^*, FRT80B”) mutant alleles. Notice the abnormal eye structure and significantly lower number of (F) white or (H) orange-colored ommatidia (highlighted by green arrows and dashed outline) corresponding to homozygous *hsp110^m1^*or *hsp110^m2^* cells, respectively, as compared to the dominant numbers of red-colored ommatidia corresponding to heterozygous cells in the same eyes.

In adult eyes of the MARCM flies, the homozygous *hsp110* mutant cells could be distinguished from surrounding heterozygous or homozygous WT cells based on their pigmentation levels. In the parental lines used to generate the MARCM clones, the eyes of adult flies carrying the WT MARCM chromosome had a red eye color due to the presence of multiple copies of transgenes with the eye color marker mini-White (Lee and Luo 1999) (Fig. 4D), while flies carrying the *hsp110^m1^* allele alone were white-eyed (Fig. 4E), as the insertion transposon *c00082* responsible for the *hsp110^m1^* allele lacks the mini-White eye marker, and the flies carrying the *hsp110^m2^* allele alone had a light-yellow eye color (Fig. 4G) because of the presence of a copy of the mini-White marker in the corresponding transposon *l(3)s64906*. Thus, in the eyes of the MARCM mosaic adult escapers, ommatidia homozygous for *hsp110^m1^* were devoid of pigmentation and were white-colored (arrows in Fig. 4F), ommatidia homozygous for *hsp110^m2^*were recognizable for their light-yellow color (arrow in Fig. 4H), both distinguishable from the surrounding, red-colored heterozygous WT cells carrying the MARCM chromosome. Notably, for non-balancer escapers carrying the mosaic clones for either the *hsp110^m1^* or *hsp110^m2^* alleles, their eyes were dominated by red-colored WT ommatidia, except for small patches of white (arrows in Fig. 4F for homozygous *hsp110^m1^* cells) or light-yellow (arrow in Fig. 4H for homozygous *hsp110^m2^* cells) ommatidia. Although large numbers of homozygous mutant *hsp110* cells were viable and proliferated during development (*i.e.,* GFP-positive cells in Fig. S3 and 4B, C), few survived into adulthood (Fig. 4F, H), suggesting that mutant *hsp110* cells were eliminated during later fly development.

To confirm this observation and bypass the prevalent lethality of *hsp110* mutant mosaic flies induced by ubiquitous *hs*-FLP, we used *eyeless-*FLP to induce mutant clones specifically in the developing eye, a tissue not essential for fly survival (Newsome, Asling, and Dickson 2000; Xu and Rubin 1993). In this mosaic study, the sister WT FRT80B chromosome carried a transgene for the White eye pigmentation marker (p{W+}) and a recessive cell lethal (*cl*) mutation (adult eye image in Fig. S4A), allowing us to distinguish homozygous *hsp110* mutant cells (e.g., genotype: *hsp110 ^m1^*/*hsp110 ^m1^*) from heterozygous WT cells (e.g., genotype: *hsp110 ^m1^* / *cl*, p{W+}) in adult mosaic eyes. In the resulting adult progeny, the eyes were composed almost entirely of red-colored WT ommatidia, with almost no white-colored mutants present except for remnants of scar tissue with a few white-colored ommatidia (arrow in Fig. S4C). Together, these findings support that Hsp110 is not required for cell proliferation during development but becomes essential for long-term cellular survival and maturation into adult structures.

The depletion of *hsp110* mutant cells in adults could be due to their inherited growth disadvantage in the absence of the Hsp110-regulated chaperone network, leading to mutant cells being outcompeted by surrounding WT cells during development. To test this hypothesis, we generated eye-specific mutant clones in a Minute background (M(3)RpS17^4^), using the elegant *eyeless*-FLP/Minute technique developed by the Dickson group (Newsome, Asling, and Dickson 2000). *RpS17^4^* is a dominant ribosomal gene mutant that causes slower cell division and developmental delay when present in heterozygotes and cell lethality when in homozygotes (Newsome, Asling, and Dickson 2000). For genes that do not affect cell proliferation or survival, when their mosaic mutant clones were induced using this system, the eyes would be mostly composed of cells homozygous for the studied mutant, with only a small number of heterozygous cells carrying the *RpS17^4^* mutant present in the background (Newsome, Asling, and Dickson 2000). When we used the *eyeless*-FLP to induce mosaic clones for either *hsp110^m1^*(adult eye image in Fig. S4B) or *hsp110^m2^*(adult eye image in Fig. S4F) alleles against the *Rps17^4^* background (adult eye image in Fig. S4D) specifically in developing eyes, the resulting adult eyes were severely deformed and significantly smaller (Fig. S4E, G), but were still mainly composed of red-colored WT cells, with only small patches of homozygous *hsp110* mutant cells present, recognizable by their white (Fig. S4E for *hsp110^m1^*) or light-yellow (Fig. S4G for *hsp110^m2^*) ommatidia colors. These data support that even when provided with a growth advantage (*i.e.,* compete against surrounding cells harboring a dominant *RpS17^4^* mutation), Hsp110-deficient cells were still actively eliminated during development.

### Hsp110 affects neuronal differentiation non-autonomously in developing eyes

Given that in the developing eye and other tissues, mosaic induction produced large clones containing tens or even hundreds of *hsp110* mutant cells (Fig. S3 and 4A-C), the absence of the mutant cells in the resulting adult flies could be due to their failure to properly differentiate, thus not being able to contribute to the final adult structures and eventually eliminated. To test this, we examined how Hsp110 affected neuronal differentiation in the eye, a well-characterized tissue, extensively documented for its orderly developmental processes and stereotypic patterning (Wolff and Ready 1991; Wolff and Ready. 1993; Kumar 2012). The waves of neuronal differentiation that contribute to the final adult eye structure initiate during the third-instar larval stage, at the posterior end (“p” in Fig 4A-C) of the eye imaginal disc where it connects to the brain lobe via the optic stalk, and gradually progress to the anterior end (“a” in Fig 4A-C), in a process that is marked by the characteristic anterior migration of the morphogenic furrow (MF) sweeping through the eye disc. Neuronal differentiation in the eye discs can be clearly visualized by staining with well-established neuronal markers such as HRP that labels neuronal membrane (Fig. 4A) (Jan and Jan 1982; Wolff and Ready 1991). Interestingly, in eye imaginal discs of MARCM larvae that harbored large mosaic clones induced by *hs*-FLP for either *hsp110^m1^* or *hsp110^m2^* mutant alleles, (Fig 4A-C), HRP staining revealed that, compared to the age-matched WT controls, which exhibited orderly recruitment of differentiated neurons into rows of maturing ommatidia along the posterior-to-anterior axis (arrows in Fig. 4A), some showed orderly recruitment of neuronal photoreceptor cells (arrows in Fig. 4C), and others exhibited little to no detectable HRP signals from differentiated neurons (Fig. 4B).

To further confirm this observation, we stained the eye imaginal discs for the glycoprotein Chaoptin, another well-established marker for the membrane of neuronal photoreceptor cells in the developing eye field and the axonal bundle from the Bolwig’s organ - the light-detecting larval eye that travels through the optic stalk into the brain (Fig. 5A) (Fujita et al. 1982). Similar to that observed by anti-HRP staining (Fig. 4A-C), eye imaginal discs that harbored large clones homozygous for either *hsp110^m1^*or *hsp110^m2^* mutant alleles contained a reduced number of Chaoptin-positive neuronal photoreceptor cells at the posterior end of the eye discs (Fig. 5B), or hardly any (Fig. 5C), as compared to those in the WT control of the same developmental stage (Fig. 5A). However, the Chaoptin-positive axonal bundle from the Bolwig’s organ was clearly visible in the eye discs of both mosaic mutants and WT controls, demonstrating that the observed differences did not arise from an issue with tissue staining. Together, these findings support that Hsp110 plays a crucial role in the proper neuronal differentiation of developing flies.

**Fig 5.**
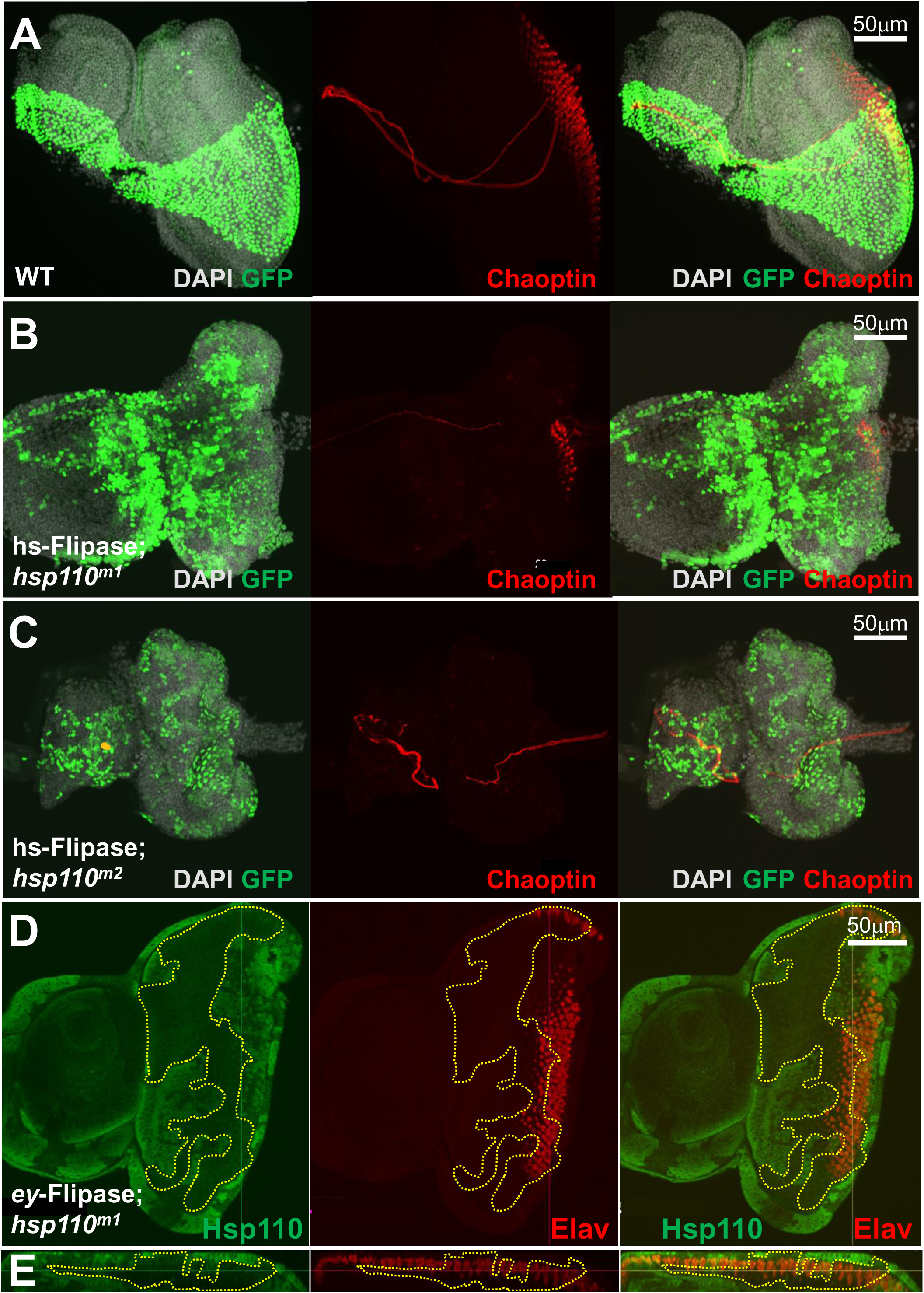
A non-cell autonomous role of Hsp110 in neuronal differentiation. Confocal images of eye imaginal discs from third-instar larvae with mosaic clones from (A) WT control, (B, D) *hsp110^m1^* or (C) *hsp110^m2^* mutant alleles, in single channels or overlaying images, as annotated. All samples are oriented with the posterior to the right and the antenna area to the left. (A-C) Eye imaginal discs with *hs*-FLP induced MARCM mosaic clones (green), co-stained against DAPI (blue) and Chaoptin (red). Notice the presence of Chaoptin-positive axonal bundles from Bolwig neurons that traverse through the eye imaginal discs. (D, E) Single-section confocal imaging, visualized from (D) X-Y or (E) X-Z dimensions, of a third-instar eye imaginal disc carrying *eyeless-*FLP-induced mosaic clones homozygous for the *hsp110^m1^* mutant allele, double-labeled for Hsp110 (green) and Elav (red). The colored lines highlight the planes of the confocal imaging. Notice the overlay of Elav-positive neurons with homozygous *hsp110^m1^ mutant* cells inside the boundary of mosaic clones, recognizable for their absence of endogenous Hsp110 (highlighted by the dashed yellow lines). Genotype: *y, w, ey-*FLP2/+; M(3)Rps17^4^, P{w+}, FRT80B/ *hsp110^m1^*, FRT80B.

In both HRP and Chaoptin staining (Fig. 4A-C and Fig. 5A-C), variable degrees of defects in neuronal differentiation were observed in the eye imaginal discs of *hsp110* mutant clones induced by *hs*-FLP, with some showing relatively normal differentiation (Fig. 4C), while others contained fewer or negligible differentiating photoreceptor cells (Fig. 4B, 5B, C). Given that in these studies, the FLP expression was under the control of the ubiquitous *heatshock* promoter, which randomly induced mosaic clones in eye imaginal discs and all other dividing tissues, such a variation could be due to differences in the sizes, timing or location of the induced mutant clones. It also raised the possibility that the neuronal differentiation defects in the eye discs were due to a non-autonomous effect of Hsp110 in other tissues.

To determine whether Hsp110 was required cell autonomously for neuronal differentiation, we used the *eyeless*-FLP/Minute technique to generate homozygous *hsp110* mutant clones exclusively in the eye imaginal discs (Newsome, Asling, and Dickson 2000). The third instar larval eye discs were co-labeled with antibodies for Elav, a nuclear protein that marks all differentiated neurons (O’Neill et al. 1994), and Hsp110, to negatively mark the clones of homozygous *hsp110* mutant cells recognizable for their loss of endogenous Hsp110 protein (Fig. 5D, E). Interestingly, in all examined eye imaginal discs, well-patterned rows of Elav-positive neurons were observed within *hsp110* mutant clones (highlighted by dashed lines in Fig. 5D, E), suggesting that cells depleted of endogenous Hsp110 could still differentiate into neuronal photoreceptor cells. Consistently, when the eye imaginal discs with *hsp110* mosaic clones were co-labeled with HRP and Chaoptin, we observed Hsp110-depleted cells positive for both HRP and Chaoptin (Fig. S5). Together, these results support that Hsp110 is not required cell-autonomously for neuronal differentiation, and in eye imaginal discs from animals with heat shock-induced mosaic clones (Fig. 4B and 5B, C), the failure in the differentiation of photoreceptor cells potentially arose due to a non-autonomous role of Hsp110 in other tissues outside the eye discs.

### Hsp110 affects glial cell migration in developing eyes non-autonomously

In developing eyes, neuronal differentiation is tightly integrated with the development of the glial network, which ensheathes and insulates axons to support their proper projection into the optic field of the brain, a process critical for the integrity of the visual system (Silies et al. 2007). When neuronal differentiation occurs in developing eye discs, glial cells, which originate from the brain and enter the eye from the connecting optic stalk (Fig S6A), always migrate behind the differentiating neurons, following their path gradually to the anterior end, but never overpassing the boundary of the MF that demarcates the line of neuronal differentiation (arrows in Fig. 6A and S6A), as revealed by co-staining with Elav for differentiation neurons, or HRP for neuronal membrane, and Repo for glial cells (Silies et al. 2007). Interestingly, when large patches of mutant clones were induced for either of the *hsp110* mutant alleles in the developing eyes by the *eyeless*-FLP/Minute system, glial cells frequently over-migrated to the front of the differentiating neurons, transgressing into the anterior of the eye discs (arrows in Fig. 6B, 6C, and Fig. S6B), a phenotype never observed in WT controls (Fig. 6A, S6A).

**Fig 6.**
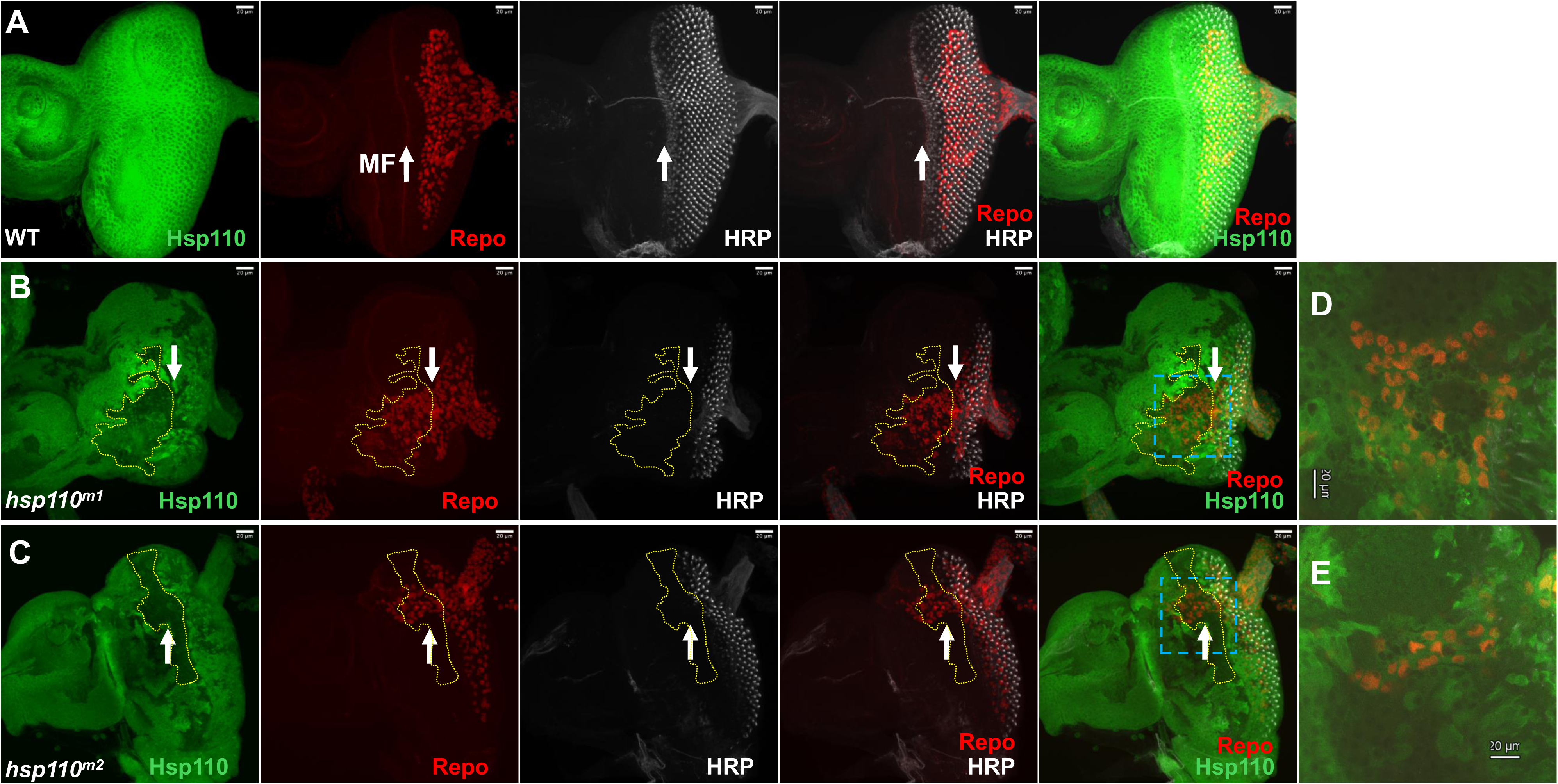
Hsp110 affects glial cell migration. Confocal images of third-instar larval eye imaginal discs from (A) WT control or (B, C) flies carrying mosaic clones for (B) *hsp110^m1^*or (C) *hsp110^m2^* mutant alleles, triple-labeled for Hsp110 (green), glial marker Repo (red), and neuronal membrane marker HRP (gray), in single channel or overlaying imaging, as annotated. Arrows highlight the boundary of neuronal differentiation along the morphogenetic furrow (MF). Dashed yellow lines marked the boundary of mosaic clones homozygous for *hsp110^m1^* or *hsp110^m2^*mutants, recognizable for their absence of endogenous Hsp110 staining signal (green). All samples are oriented with the posterior to the right and the antenna area to the left. (D, E) High magnification views of the regions highlighted within the yellow dashed lines in (B) and (C), respectively. Genotypes: (A) *w^1118^*. (B) *y, w, eyFLP2*/+; M(3)Rps17^4^, P{w+}, FRT80B/ *hsp110^m1^*, FRT80B. (C) *y, w, eyFLP2*/+; M(3)Rps17^4^, P{w+}, FRT80B/ *hsp110^m2^*, FRT80B.

Migration and differentiation of glial cells in eye imaginal discs are processes regulated by signals from the differentiating neuronal photoreceptor cells. The differentiating photoreceptor cells provide a permission signal allowing glial cells into the eye imaginal discs and an inhibitory signal that stops them just behind the MF and along the line of differentiating neurons to initiate their differentiation into wrapping glia (Franzdottir et al. 2009; Silies et al. 2007). Notably, the glial over-migration phenotype occurred mostly in regions with large patches of mutant *hsp110* cells (Fig. 6B, C). The *eyeless*-FLP system restricts the induction of *hsp110* mutant clones to cells derived from the eye imaginal disc, including the differentiated neuronal photoreceptor cells, but does not affect the brain-derived glial cells. Indeed, the over-migrated glial cells showed normal levels of endogenous Hsp110, surrounded by cells depleted of Hsp110, as revealed by co-labeling with antibodies against Hsp110 and glial cell marker Repo (Fig. 6D, E). These results suggest that the glial over-migration phenotype is not due to a loss of Hsp110 in glial cells, but rather to its absence in surrounding cells of the imaginal disc origin, most likely the differentiating neurons that normally produce the inhibitory signal to restrict the migration of glial cells.

### Higher levels of Hsp110 are detrimental to *Drosophila*

Considering that Hsp110 has been isolated as a strong protein aggregate suppressor, a potential toxicity modifier in several protein misfolding disease models, and its implication in resistance to AD in the human population, increasing the expression of the chaperone might have cytoprotective effects. However, in a previous study we observed that overexpression of Hsp110 in developing eyes caused abnormalities in the internal structures of adult eyes (Zhang et al. 2010). A fly compound eye is composed of about 800 ommatidium units, and each ommatidium contains eight stereotypically patterned photoreceptor cells recognizable for their characteristic, round-shaped rhabdomeres (Wolff and Ready. 1993), the light-detecting structure packed with actin and photon-sensitive rhodopsins, the G-protein-coupled receptors (GPCRs) that mediate phototransduction. To confirm the earlier observation, we directed Hsp110 overexpression from UAS-based transgenes specifically in the fly eye. We used the GMR-GAL4 driver to intiate a high level of transgene expression that would continue into adulthood, in all cells posterior to the MF of the eye imaginal discs from third instar larvae (Ellis, O’Neill, and Rubin 1993; Hay, Wolff, and Rubin 1994). On cross sections of control adult eyes overexpressing a transgene for UAS-luciferase dsRNA (LucdsRNA as control) (Fig. 7A), repetitive units of seven rhabdomeres in trapezoid shape were robustly visible by F-actin staining in each section layer (Fig. 7E, with Fig. 7I showing the quantification of rhabdomere numbers in analyzed eye sections), same as in WT (Wolff and Ready. 1993). However, in Hsp110 overexpressing flies, patterning of internal ommatidia was severely disrupted, with some rhabdomeres becoming abnormally elongated, others significantly smaller and fragmented, and the number of rhabdomeres varying among each ommatidium unit (Fig. 7F and quantified in Fig. 7I). Although the internal disruption was extensive, their external eye morphology appeared normal (Fig. 7B).

**Fig 7.**
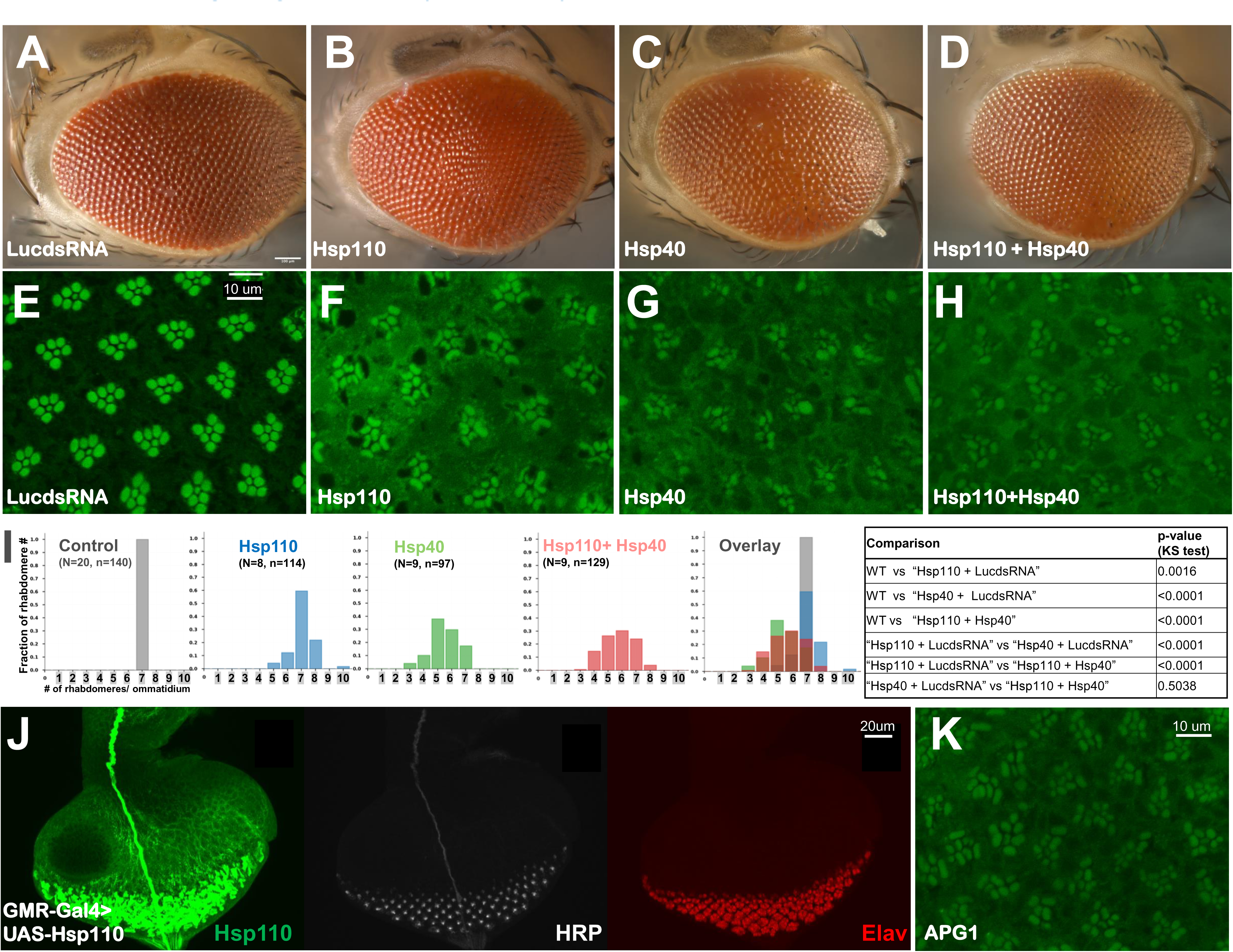
High levels of Hsp110 and Hsp40 are detrimental *in vivo*. (A-D) Bright-field imaging of external eye morphology and (E-H and K) confocal imaging of internal ommatidia structures as revealed by phalloidin staining from adult flies with eye-specific overexpression of the following UAS transgenes: (A, E) control of Luciferase-dsRNA (LucdsRNA); (B, F) Hsp110 together with LucdsRNA, (C, G) Hsp40 together with LucdsRNA, (D, H) Hsp110 together with Hsp40, (K) human APG1, all driven by GMR-GAL4. (I) Bar graph presentation on the distribution patterns on the fraction (Y-axis) of the numbers of rhabdomere per ommatidium (X-axis) in the examined genotypes from (E-H), respectively, with a table summary of the statistical difference between the studied genotypes, as annotated. N: number of independent eye samples quantified. n: number of total rhabdomeres quantified in each genotype. (J) Confocal imaging of an eye imaginal disc from third-instar larvae with eye-specific overexpression of Hsp110 driven by GMR-GAL4 (genotype: GMR-Gal4/+; UAS-Hsp110/+), triple-labeled for Hsp110 (green), HRP (gray), and Elav (red), as annotated. Notice the high levels of expression of Hsp110 in all cells posterior to the morphogenetic furrow. The eye disc is oriented with the anterior at the top and the posterior at the bottom.

To examine whether the phenotype in Hsp110 overexpressing flies was due to a disruption of neuronal differentiation during earlier eye development, or arose in later maturation stages, we co-stained the third larval eye imaginal discs for Hsp110 and neuronal markers HRP and Elav. The staining revealed that despite significantly high levels of Hsp110 in cells post the MF, differentiation and patterning proceeded normally, with stereotypical, progressive recruitment of differentiated neurons into maturing ommatidial clusters along the posterior to anterior axis of the eye imaginal discs (Fig. 7J). Together, they suggest that abnormally high levels of Hsp110 do not affect eye differentiation processes but potentially disrupt rhabdomere growth and patterning during later maturation.

To test whether the detrimental effect was limited to the eye, we targeted Hsp110 overexpression ubiquitously in all tissues, driven by tubulin-GAL4. Importantly, in four independent UAS-Hsp110 transgenic lines we tested, high levels of Hsp110 expression all caused animal lethality, with most of the Hsp110-overexpressing flies dead around the pupal stage. Together these findings demonstrate that high levels of Hsp110 can be detrimental in different tissues, and that both its levels and activity need to be properly modulated to support normal animal development and physiology.

### Higher levels of Hsp70 co-chaperone Hsp40 are similarly detrimental to Drosophila

In the current model of the Hsp70 chaperone cycle, Hsp110 coordinates with Hsp40 to facilitate Hsp70 transition through the successive steps of ATP-dependent substrate binding, unfolding, release, and refolding. Thus, engineered higher levels of Hsp110 might disrupt Hsp70 regulation, leading to an unproductive chaperone cycle and subsequent secondary effects such as proteostatic stress. Such a scenario would predict that abnormally high levels of Hsp40, the other key regulator of the Hsp70 chaperone cycle, would cause a similar detrimental effect as Hsp110. Indeed, when ectopically expressed in developing eyes by GMR-GAL4, Hsp40 also induced similar rhabdomere structure and organizational phenotypes as those caused by excess Hsp110, despite apparently normal external eye morphology (Fig. 7C). In addition to abnormally elongated or fragmented rhabdomeres, missing rhabdomeres and significant disruption of the stereotypical trapezoid patterning were observed (Fig. 7G).

The above hypothesis also raises the possibility that simultaneously increasing the levels of both Hsp110 and Hsp40 to match stoichiometry might rebalance the equilibrium of the Hsp70 chaperone cycle, lessening the detrimental effects. Alternatively, if the detrimental effects are due to the excessive, independent activities of Hsp110 and Hsp40 unrelated to the Hsp70 chaperone cycle, then co-expression of Hsp110 with Hsp40 should lead to an additive, more severe consequence. To distinguish between these two possibilities, we examined fly eyes with simultaneous co-expression of Hsp110 and Hsp40 under the GMR-GAL4 driver. Notably, the external eye morphology of such co-expression flies appeared normal (Fig. 7D), similar to Hsp110- or Hsp40-single overexpression flies (Fig. 7B, C). Internally, the ommatidia structure remained disrupted, with missing and fragmented rhabdomeres and misaligned ommatidium units (Fig. 7H).

To quantitatively evaluate the severity of the eye phenotypes induced by Hsp110 and Hsp40 overexpression alone or in combination, and to minimize the dilution effect of the GAL4 driver between flies carrying different copies of UAS transgenes, we included the UAS-LucdsRNA transgene as an internal control and quantified the number of recognizable rhabdomeres in the eyes of flies co-expressing Hsp110 with LucdsRNA, Hsp40 with LucdsRNA, and Hsp110 with Hsp40 (Fig 7I). This analysis revealed that Hsp40 overexpression caused a more severe disruption than Hsp110 (Table in Fig 7I. p< 0.0001 KS test between “Hsp110 with LucdsRNA” and “Hsp40 with LucdsRNA” groups), but their co-expression did not further enhance the phenotype, as the distorted rhabdomere distribution pattern was indistinguishable between “Hsp110 with Hsp40” and “Hsp40 with LucdsRNA” groups (Table in Fig 7I. p=0.50, KS test). Although it was difficult to quantify rhabdomere morphology, we qualitatively assessed that the overall morphology of the rhabdomeres in “Hsp110 with Hsp40” eyes (Fig. 7H) appeared less deformed than that in “Hsp40 with LucdsRNA” eyes (Fig. 7G). Together, these data align with the hypothesis that the detrimental effects of Hsp110 and Hsp40 overexpression were potentially due to their shared roles in Hsp70 regulation, and the physiological levels of Hsp110 and Hsp40 co-chaperones need to be properly balanced for Hsp70 to efficiently execute a productive unfolding/refolding cycle.

Lastly, given the significant sequence and structural conservation between fly and human Hsp110 proteins, we tested the overexpression effect of transgenic human Hsp110 in the fly eye. Remarkably, when directed by the GMR-GAL4 driver, ectopic expression of HSPA4L (APG-1) caused similar, albeit weaker, eye phenotypes as fly Hsp110 (Fig. 7K), thus supporting the functional conservation of Hsp110 from flies to humans, in what must be a fundamental and shared process. These results strongly suggest that in humans, the levels and activities of Hsp110 similarly need to be carefully modulated to ensure a productive Hsp70 chaperone cycle to support proper neural development.

## Discussion

As the first systematic study on *Drosophila* Hsp110, we demonstrated a high degree of conservation between fly and human Hsp110 proteins at the sequence, structural, and potentially functional levels. Consistent with this conservation, human APG-1 induced similar overexpression phenotypes as fly Hsp110 (Fig. 7K), and human HSPA4 and HSPA4L partially rescued the germ cell development defects of flies with RNAi-mediated Hsp110 depletion in male germlines (Houston et al. 2025). Given that a single Hsp110 gene exists in the *Drosophila* genome and considering the abundant and sophisticated experimental tools available in this prime animal model, *Drosophila* proved to be an ideal model to dissect the physiological functions and regulation of Hsp110 in metazoans.

Our results further revealed a diverse and essential role of Hsp110 in various tissues and developmental stages of *Drosophila*, including its effects on neuronal differentiation, glial cell migration, in ovary development, as well as its critical requirement for long-term cell survival. Notably, despite being isolated as a potent suppressor of protein aggregation and neurotoxicity in multiple neurodegenerative disease models, and its intricate link to AD resistance, higher levels of Hsp110 proved to be detrimental in flies, an effect mimicked by Hsp40 overexpression. However, their co-expression did not worsen the consequence. Together, these results suggest that *in vivo*, the levels and activities of Hsp110 and Hsp40 co-chaperones need to be properly balanced to achieve an efficient and productive Hsp70 chaperone cycle. Therefore, the Hsp70 chaperone network needs to be considered as a whole when targeted for therapeutic purposes. Considering the essential yet dosage-sensitive roles of Hsp110 in maintaining normal cellular physiology and survival, these results suggest the existence of robust *in vivo* regulatory mechanisms that modulate the expression and activity of Hsp110, thereby satisfying complex physiological demands under both normal and stressed conditions in multicellular organisms.

### Significant conservation of Hsp110 genes from flies to humans

Several notable similarities exist between *Drosophila* and mammalian Hsp110 chaperones, including their gene structure, amino acid sequences, conformation, and the subcellular localization of their protein products. Noticeably, both mammalian Hsp105A (HSPH1) and fly Hsp110 genes similarly encode two protein isoforms due to alternative splicing of a small coding exon for additional 32 (fly) or 44 (human) amino acid-long sequences that are located within the same unstructured loop in the SBDβ subdomain of the respective proteins (Fig. 1B). Such a conservation at the gene structure, splicing and conformational levels implies a functional importance of this splicing regulation, and potentially differential roles for the two Hsp110 isoforms at physiological levels. An earlier biochemical study demonstrated that the loop insertion in APG2 plays an important regulatory role in the Hsp70 chaperone cycle, acting as a molecular switch to facilitate the release of APG2 from the Hsp70 chaperone, thereby preventing their unproductive association (Cabrera et al. 2019). It is intriguing to speculate whether the extra sequence encoded by the alternatively spliced exon confers another layer of regulation on the physical association between Hsp110 and Hsp70.

In *Drosophila,* endogenous Hsp110 was ubiquitously present in all examined cells at various developmental stages (Fig. 3), with the shorter Hsp110-A isoform as the dominant product and its overall endogenous levels about six-fold higher than the longer Hsp110-G isoform (Fig. 2A, B). At subcellular levels, *Drosophila* Hsp110 was predominantly a cytoplasmic protein in most cells but also detected at low levels in the nucleus of neurons and glial cells, with prominent nuclear enrichment in polyploid salivary gland cells, suggesting a cell-type dependent subcellular localization pattern (Fig. 3). Interestingly, mammalian Hsp105 exhibits discrete subcellular localization in an isoform-dependent manner. The longer Hsp105A localizes mainly to the cytoplasm, is constitutively expressed, and its expression can be further induced by stresses. In contrast, the shorter Hsp105B localizes to the nucleus, and its expression is induced specifically during mild heat shock stress (Saito, Yamagishi, and Hatayama 2007, 2009). In our study, heat shock stress did not significantly increase the overall level of endogenous Hsp110 protein in *Drosophila* (Fig. 2A, B). Currently, the available tools do not allow us to determine whether the discrete localization patterns of fly Hsp110 are also isoform-specific, as has been observed for mammalian Hsp105 isoforms. It will be important to determine how this differential subcellular localization of Hsp110 proteins is controlled, and its physiological significance.

In addition to cytoplasmic and nuclear localization, Hsp110 was also observed in neuromuscular junctions (Fig. 3C), near or on the plasma membrane (Fig. 3D), and on the axonal bundle associated with the brain (Fig. 3E), a pattern especially prominent in neurons overexpressing Hsp110 (Fig. 7J). Interestingly, at high magnification, endogenous Hsp110 appeared to be enriched in minute puncta throughout the cells (Fig. 3G), raising the intriguing possibility that endogenous Hsp110 might nucleate with its chaperone partners on subcellular nanodomains to more efficiently carry out their chaperone function. Notably, a very recent study showed that both *in vitro* and in human cell lines, a chaperone protein disulfide-isomerase A6 (PDIA6) could recruit additional chaperones, including Hsp70 and Hsp40, into phase-separated condensates in the endoplasmic reticulum (ER) to promote their cooperation and efficient client processing (Leder et al. 2025). It will be important to determine if a similar mechanism is responsible for the nucleated subcellular distribution of Hsp110 in these studies.

### The essential roles of Hsp110 in animal development

Consistent with its central role in the chaperone cycle, Hsp110 is essential for fly development, with homozygous mutants dying during embryogenesis. Interestingly, mosaic analyses revealed that large mosaic clones composed of tens or even hundreds of homozygous *hsp110* mutant cells could be generated in different developing tissues, such as the brain, wing, and eye imaginal discs (Fig. S3 and Fig. 4A-C). An observation that demonstrated cells depleted of endogenous Hsp110 were not immediately eliminated, but could still survive, proliferate, and even enter neuronal differentiation during development. Given that theoretically each mosaic clone originates from a single dividing parental cell (Chou and Perrimon 1992; Xu and Rubin 1993), these findings suggest that Hsp110 is not necessary for cell proliferation. Nevertheless, mosaic analyses in developing eyes, which have highly stereotypical patterning and abundant markers to trace the fate of mutant cells, revealed that although *hsp110* mutant cells could proliferate and even differentiate into photoreceptor neurons, they were eventually eliminated and rarely contributed to adult structures (Fig. 4 and Fig. S4), thus revealing the critical role of Hsp110 for long-term cell survival. These findings raise more questions that need to be addressed in future studies, such as what activates this Hsp110-associated cell elimination process, if cell death is due to disrupted proteostasis and subsequently heightened cell stress, and whether it is executed through apoptosis or other mechanisms.

Mosaic analyses also revealed a strong effect of Hsp110 on neuronal differentiation, as eye imaginal discs containing large mutant clones showed notably reduced numbers (Fig. 5B and Fig. 6), or even a complete absence (Fig. 4B and Fig. 5C) of differentiating neurons, a phenotype never observed in WT animals. Further analyses suggest that this neuronal defect might be mediated partially through a non-autonomous role by Hsp110 during development, as the most severe phenotype (*i.e.,* an absence of neuronal differentiation in the developing eyes, Fig. 4B and Fig. 5C) was observed only when *hsp110* mutant clones were induced ubiquitously by *hs*-FLP, which is under the control of a *heatshock* promoter that targets most dividing tissues, not just the eyes. When the mutant clones were specifically restricted to the eye imaginal discs by eye-specific *eyeless*-FLP, a prominent number of differentiated neurons homozygous for *hsp110* mutants were observed (Figs. 5D, E, and S5, Fig. 6). These results raise the possibility that loss of Hsp110 affects tissue-tissue or cell-cell communication, and the failure of neuronal differentiation in *hsp110* mosaic mutants was likely due to a role of Hsp110 in other non-eye tissues that normally provide permission, or instruction signals to initiate neuronal differentiation in the developing eye discs. One such potential candidate is the steroid hormone ecdysone from the endocrine prothoracic gland. The ecdysone surge during late third instar larval stage is necessary for the initiation of eye differentiation and the subsequent orchestration of multiple local signaling pathways establishes the stereotypical patterning of photoreceptor cell differentiation (Treisman and Heberlein 1998; Brennan, Ashburner, and Moses 1998; Kumar 2012). If Hsp110 is required in the ecdysone-producing prothoracic gland, then its depletion might dampen the ecdysone surge, blunting neuronal differentiation in the eye discs.

A similar scenario may also apply to the glia cell over-migration phenotype in the eye imaginal discs, which contain large mosaic clones for *hsp110* mutants (Fig. 6, Fig. S6). In this study, WT glia cells originating from the brain trespassed beyond the frontline of differentiating neurons, a phenomenon that never occurs in normal situations (Fig. 6, Fig. S6). As glial cell migration is normally restrained by the neuron-derived FGF8-like ligand, which binds to and activates the fibroblast growth factor (FGF) receptor on glial cells to instruct their fate switch from migration to differentiation (Franzdottir et al. 2009; Silies et al. 2007), the over-migration phenotype similarly raises the intriguing possibility that Hsp110 is normally required in differentiated neurons for the production of inhibitory FGF8 ligand. Future studies are needed to test these hypotheses and determine whether Hsp110 affects specific signaling pathways such as FGF and nuclear hormone receptors, or if the observed defects are due to secondary effects from stressed *hsp110* mutant cells.

### A delicate role of Hsp110 in the chaperone cycle and cellular homeostasis

Considering the growing link between Hsp110-regulated chaperone machinery and a diverse group of protein misfolding diseases, it is critical to understand how innate cellular protective mechanism can be employed for disease prevention and treatment. Our results suggest a delicate, double-edged role of Hsp110 in animal physiology and neuronal function, as both its depletion and aberrant elevation cause detrimental effects to neurons and the animal as a whole. A similar double-edged effect was also reported in C. *elegans,* as both RNAi-mediated depletion or overexpression of the C. *elegans* Hsp110 gene lead to reduced lifespan and low fecundity in the worm (Montresor et al. 2025), supporting a conserved, dichotomous role of Hsp110 in cellular health and animal fitness. Given that overexpression of Hsp40 causes similar detrimental effects (Fig. 7G), and that their co-expression does not further enhance the consequence (Fig. 7H, I), the detrimental effect by Hsp110 and Hsp40 is likely due to disruption of the Hsp70 chaperone cycle. Importantly, existing biochemical evidence supports this hypothesis. For example, excessive Hsp110 inhibits the ATPase and chaperone activity of HSP70 (Rampelt, Mayer, and Bukau 2018), and higher concentrations of APG2 inhibit, instead of promoting, the disassembly of luciferase aggregates (Aguado et al. 2015). Similarly, excess equimolar amounts of yeast Sse1 block the unfolding/refolding cycle of denatured MlucV by Hsp70 protein Ssa1, leading to an increased aggregation of misfolded MlucV (Rebeaud et al. 2025).

The above findings also raise questions concerning Hsp110 regulation, both in terms of its expression and activity, as well as its physical association with Hsp70 and other potential partners *in vivo*. Although a detailed understanding on the regulatory mechanisms of metazoan Hsp110 is still lacking, recent biochemical studies by the Moro group showed that the loop insertion in the SBDβ subdomain promoted efficient disassembly of a APG2/HSC70 complex, while in the C-terminal extension of APG2, the α-helical region interacted with the NBD domain of Hsp70 in an ATP-dependent manner to promote the stability of the APG2/HSC70 complex, and its IDR subdomain downregulated nucleotide exchange activity for HSC70 (Cabrera et al. 2022; Cabrera et al. 2019). Together these findings suggest potentially important roles of the two largely unstructured extensions in Hsp110 regulation. Notably, these two extensions are markedly elongated in metazoan Hsp110 as compared to those from single-cell species. Furthermore, the loop insertion has two alternatively-spliced forms that are conserved from flies to humans, while the C-terminal extension contains many potential or validated post-translational modification (PTM) sites (Akimov et al. 2018; Cabrera et al. 2022; Velasco et al. 2019; Wagner et al. 2011), raising an attractive possibility that both the alternative splicing and the PTMs serve as important regulatory targets by upstream regulators to fine-tune the activity of Hsp110 and optimize the efficacy of productive Hsp70 chaperone cycles. Future challenges include, understanding the functional importance of these conserved metazoan-specific motifs and PTMs under physiological and pathological settings, especially in long-lived cells such as neurons.

## Supplemental Figure legends

**Supplemental Figure S1.**
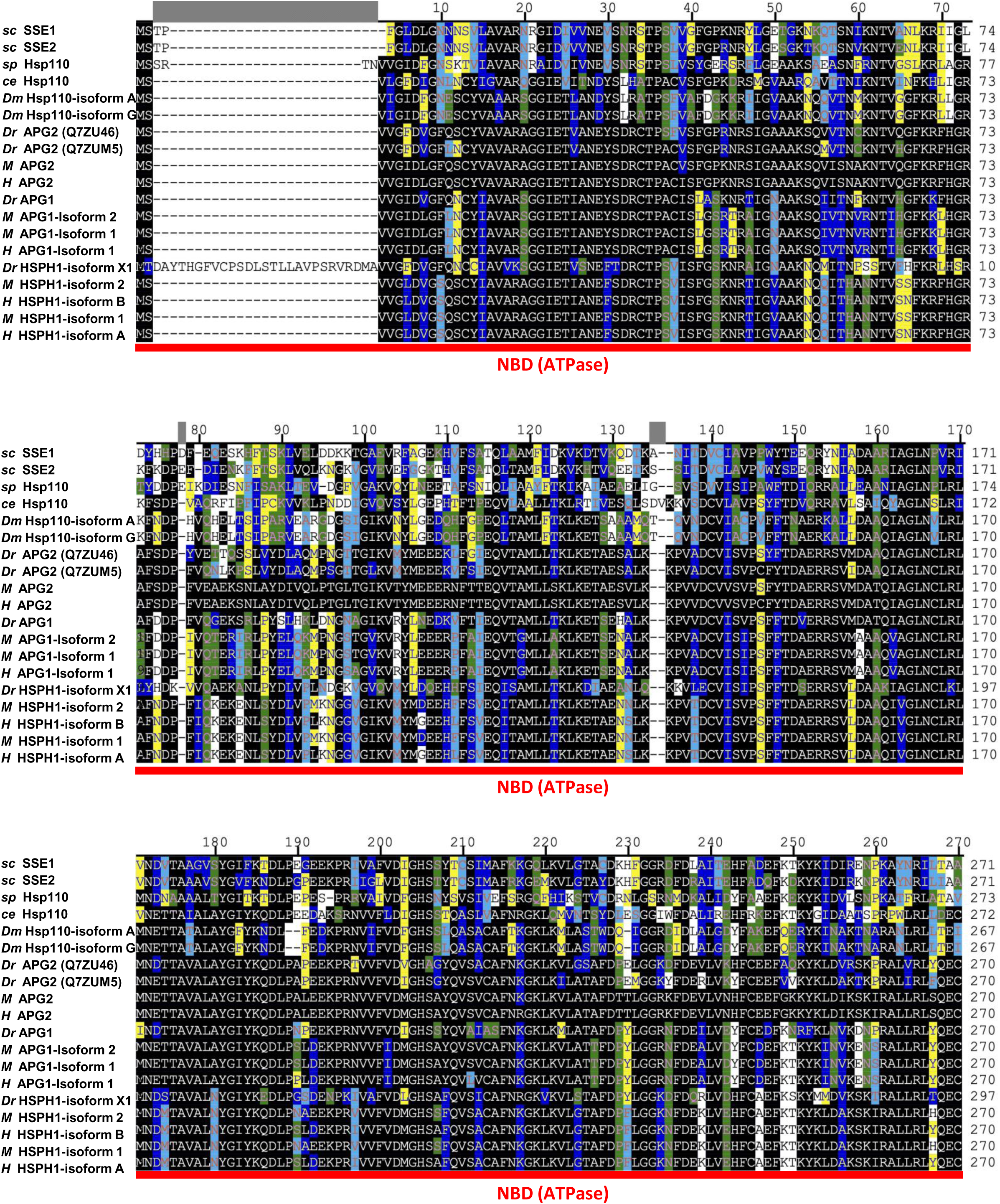
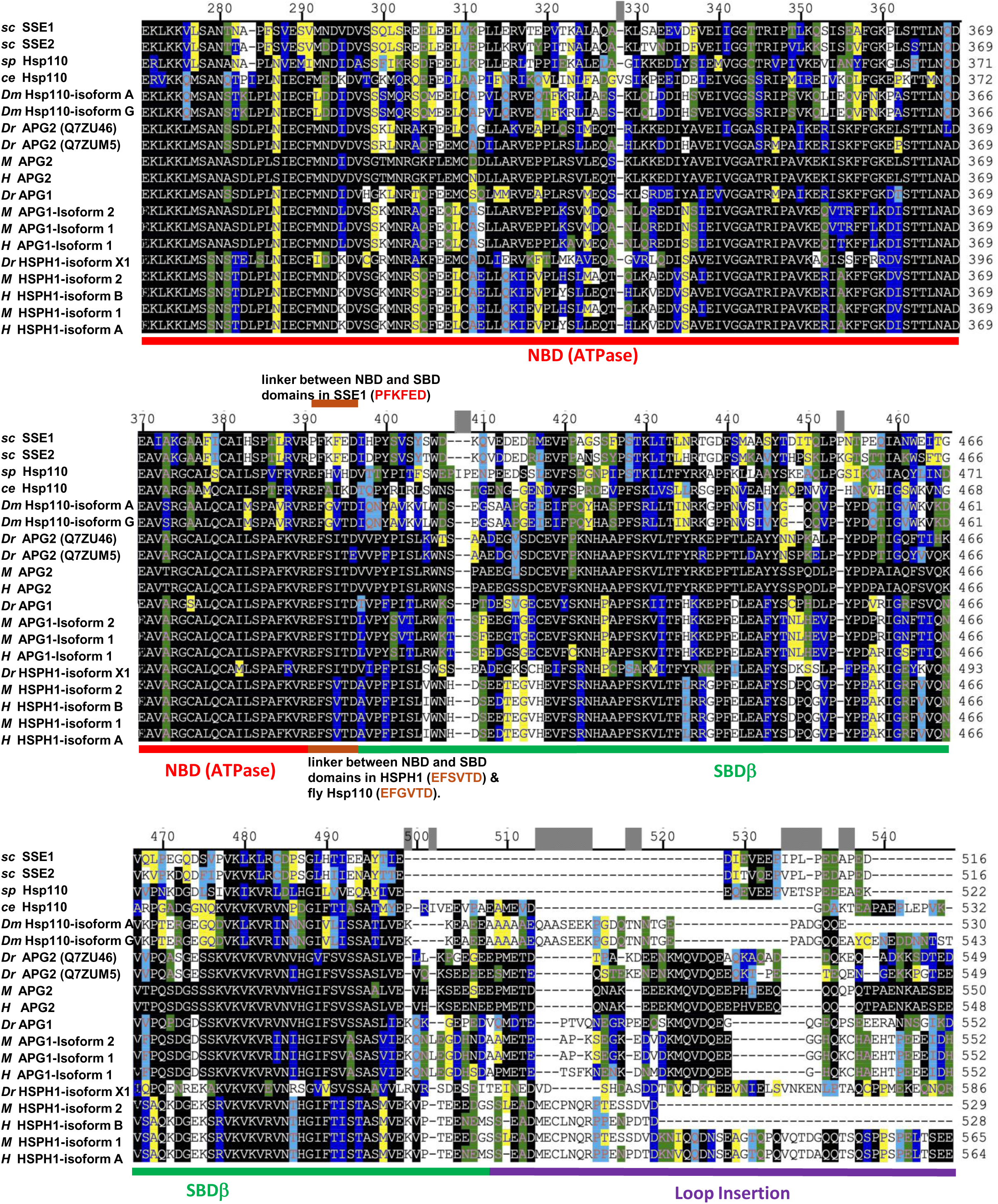
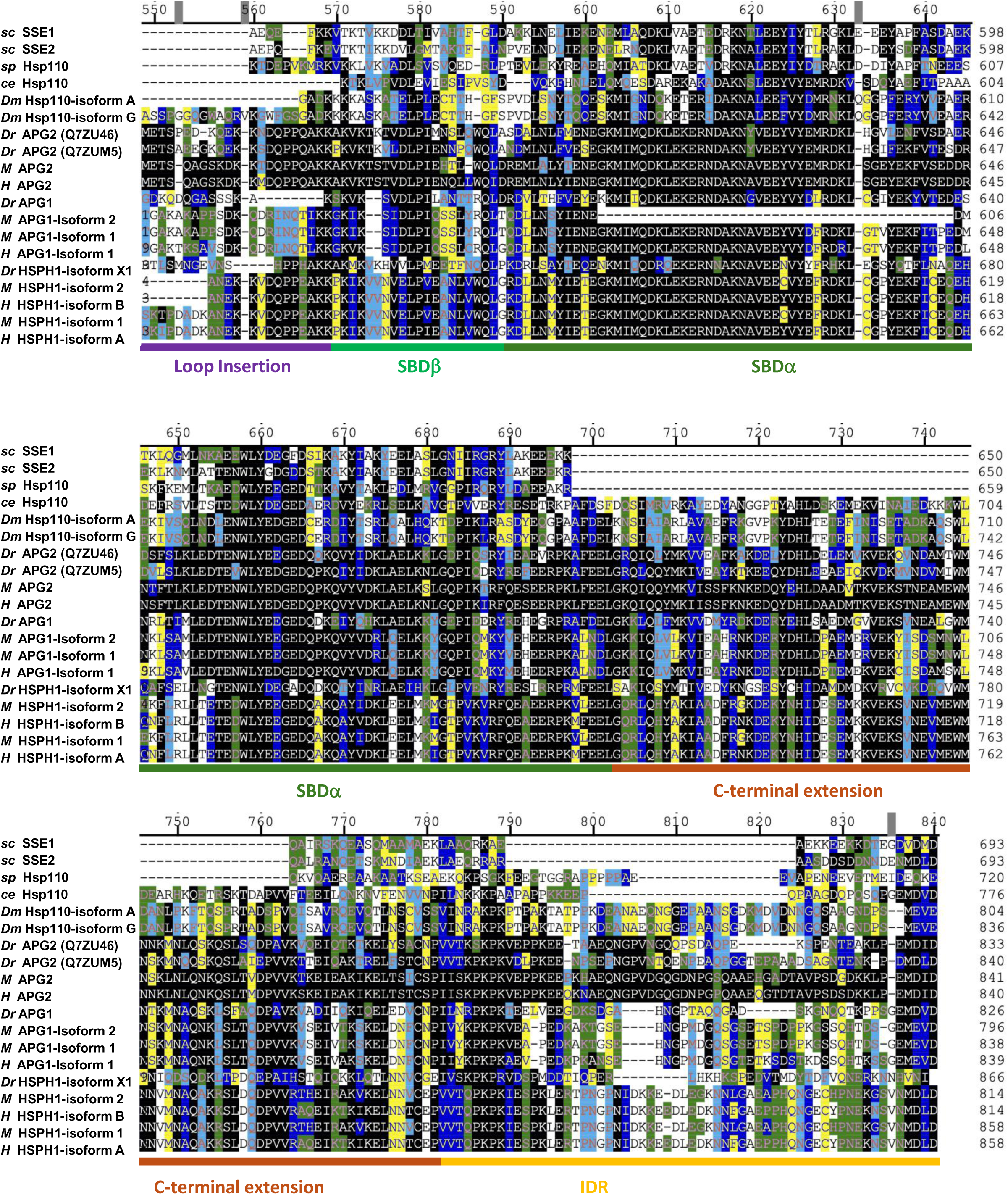
Conservation of Hsp110 proteins during evolution. Sequence alignment of Hsp110 from representative species, as annotated, aligned using the Cluster V method. The following color decorations in the alignment indicate the properties of amino acid similarities, colored with the following hierarchy order (from low to high): Cyan: Charge. Acidic (D, E), basic (H, K, R), neutral (A,C,F,G,I,L,M,N,P,Q,S,T,V,W,Y). Green: Structural similarity. Ambivalent (A,C,G,P,S,T,W,Y), external (D,E,H,K,N,Q,R), internal (F,I,L,M,V) Yellow: functional group. Acidic (D,E), basic (H,K,R), f-hydrophobic (A,F,I,L,M,P,V,W), p-polar (C,G,N,Q,S,T,Y). Blue: chemical group. Acidic (D,E), basic (H,K,R), aliphatic (A,G,I,L,V), amide (N,Q), aromatic (F,W,Y), hydroxyl (S,T), amino (P), sulphur (C,M)). Black decorates amino acids that are identical to the sequence of human APG2 (hHspA4-P349312) in the alignment. The scale of the alignment is drawn according to the sequence of human APG2 (hHspA4-P349312).

**Fig S2.**
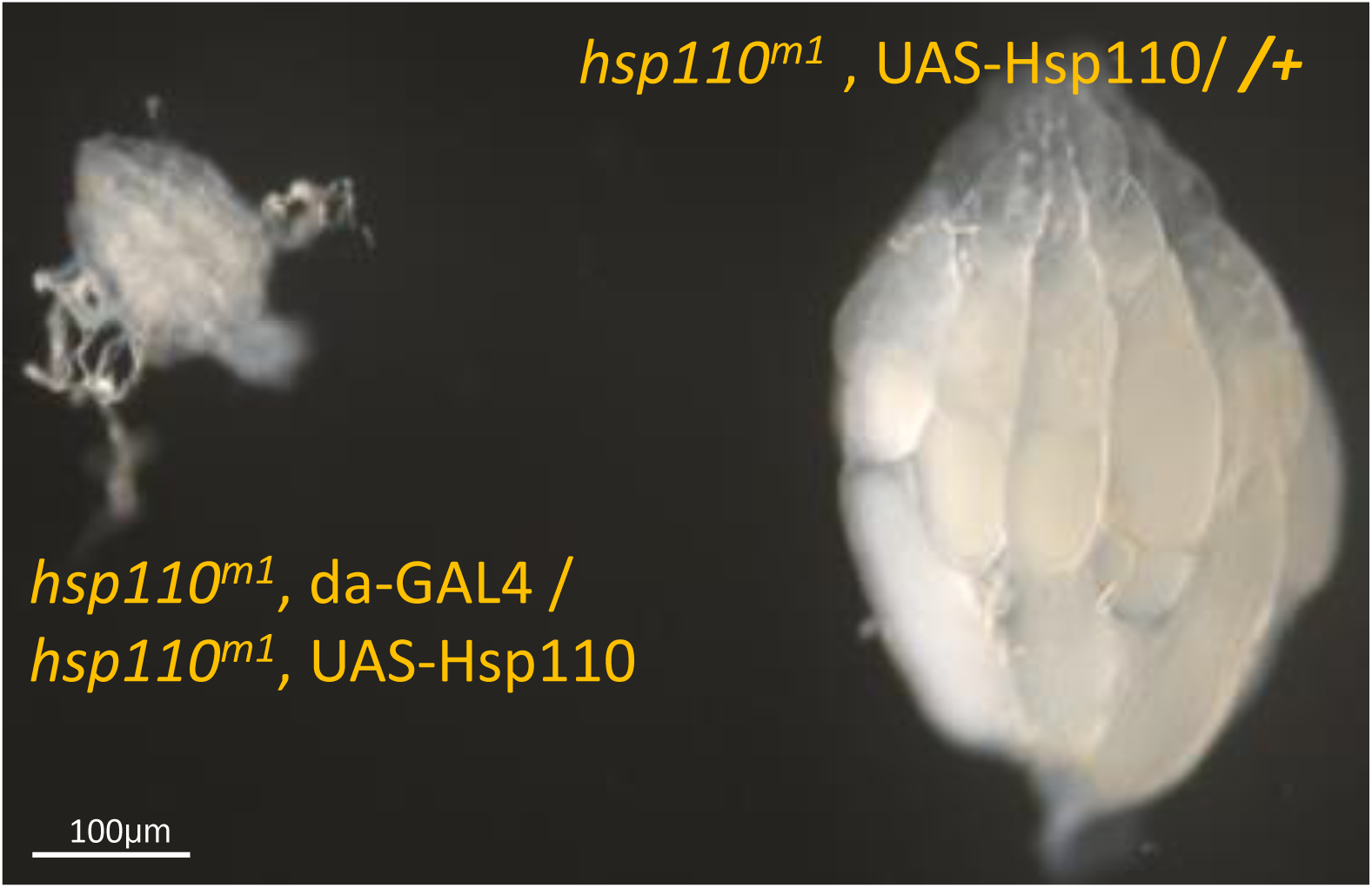
Hsp110 is essential for ovarian development. Stunted ovary development in a homozygous *hsp110^m1^* mutant adult female rescued by ectopically expressed Hsp110-A isoform from UAST-*hsp110* transgene driven by da-GAL4 (genotype: *hsp110^m1^*, da-GAL4 / *hsp110^m1^*, UAS-*hsp110*). Notice the significantly smaller ovary from the rescued female as compared to the age-matched control of a heterozygote *hsp110^m1^* female (genotype: *+* / *hsp110^m1^*, UAS-*hsp110*). Female flies were fed with yeast for 24 hours before the dissection.

**Fig S3.**
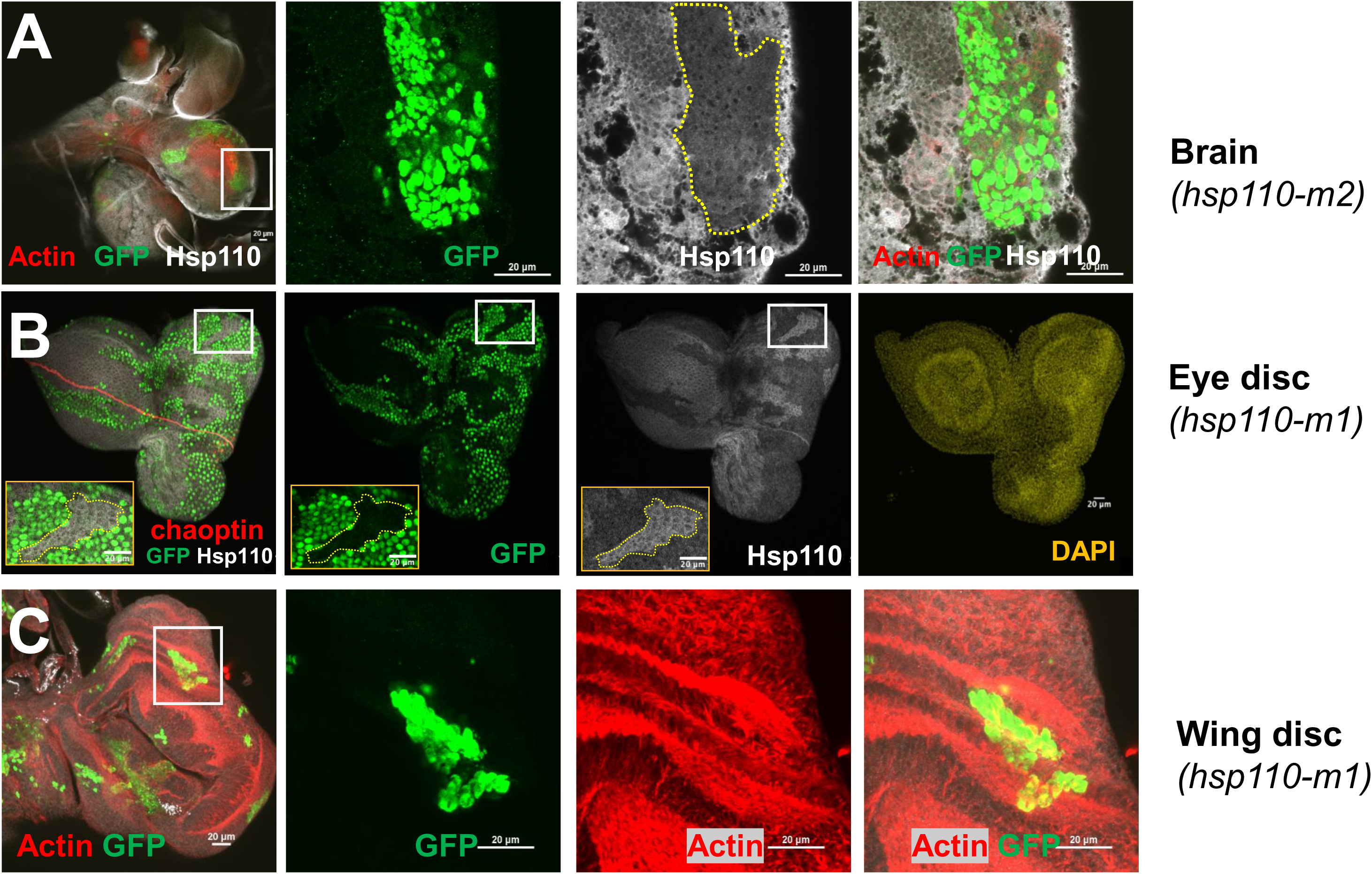
Hsp110 is dispensable for cell proliferation. Confocal images of mosaic clones homozygous for *hsp110^m1^* mutant cells (green) generated by the heat shock-induced MARCM system in (A) a brain co-stained for Hsp110 (gray) and F-Actin by phalloidin, (B) eye imaginal disc co-stained with Hsp110 (gray), and DAPI (yellow), (C) wing imaginal disc co-stained for F-Actin (red), all from third-instar larvae, shown in single channels or overlaying images, as annotated. White boxes highlight the regions shown in the zoomed-in view of each tissue, which in (B) are indicated by yellow insets. Yellow dashed lines in (A, B) highlight the boundary of mutant clones that show reduced levels of endogenous Hsp110 in GFP-positive mutant clones.

**Fig S4.**
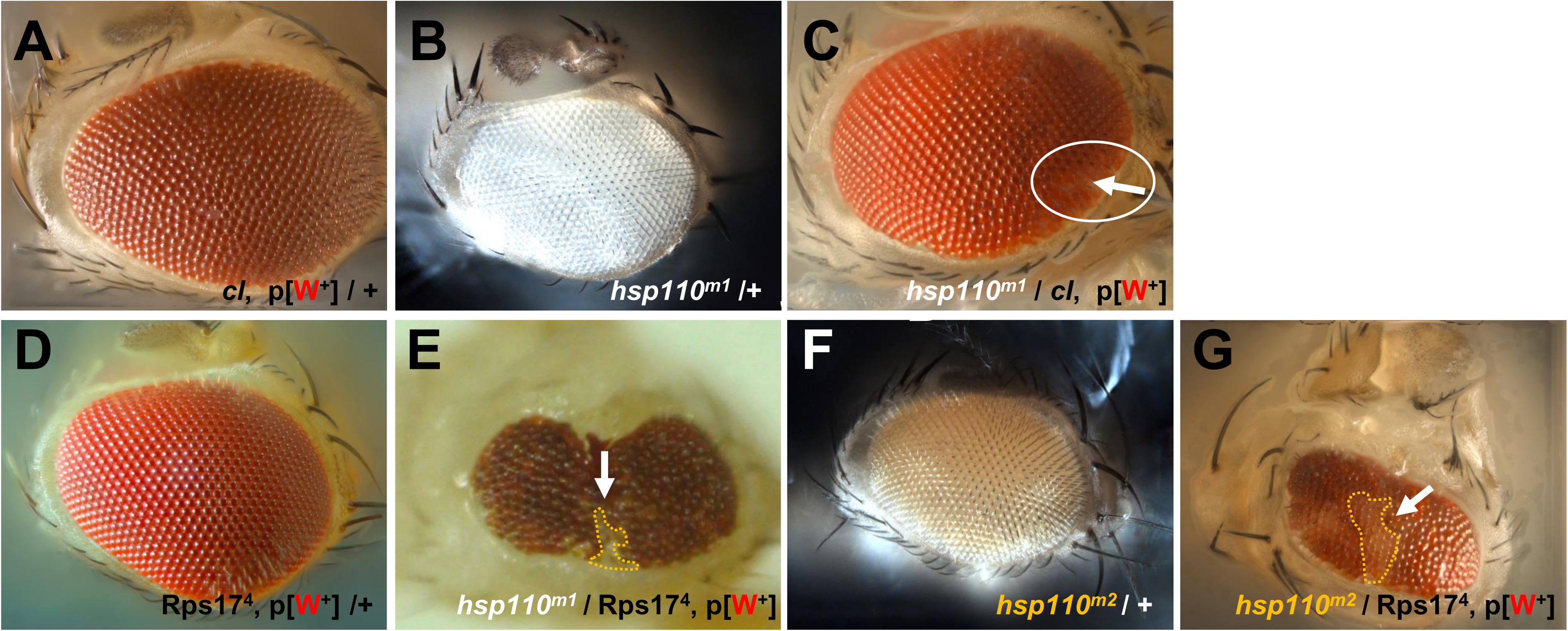
Hsp110 is required for long-term cell survival. Bright-field images of adult eyes. (A) Control of the parental line carrying the WT FRT80B chromosome marked by the transgene for eye pigmentation gene White, which produces bright red eye pigmentation (genotype: *y, w, ey-FLP2/+; cl*, P{w+}, FRT80B/+). (B, F) Controls of the parental heterozygous WT over (B) *hsp110^m1^*-FRT80B (genotype: *y, w*; *hsp110^m1^*, FRT80B/ +) or (F) *hsp110^2^*-FRT80B (genotype: *y, w*; *hsp110^m2^*, FRT80B/ +) mutant lines with white or orange eye color, respectively, due to the absence or presence of the eye pigmentation gene White in its genome. (C) A representative eye with mutant clones for *hsp110^m1^* generated by *eyeless-*FLP (genotype: *y, w, ey-FLP2/+; cl,* P{w+}, FRT80B/ *hsp110^m1^*, FRT80B). Notice the scar containing a few homozygous *hsp110^m1^* mutant cells, recognizable for their white-colored ommatidium (highlighted by the white arrow within the cycled area) in the eye. (D) Control of the parental line for WT FRT80B chromosome carrying the Minute M(3)RpS17^4^ mutation and the transgene for eye pigmentation marker White (genotype: *y, w, eyFLP2/+;* M(3)RpS17^4^, P{w+}, FRT80B/ +). (E, G) Mosaic eyes in Minute background with *eyeless*-FLP induced clones homozygous for (E) *hsp110^m1^* (genotype: y, w, eyFLP2/+; M(3)RpS17^4^, P{w+}, FRT80B/ *hsp110^m1^*, FRT80B), or (G) *hsp110^m2^* (genotype: y, w, eyFLP2/+; M(3)RpS17^4^, P{w+}, FRT80B/ *hsp110^m2^*, FRT80B) mutants, recognizable for their white or orange ommatidium colors (highlighted by white arrows and outlined with yellow dashed lines), respectively, surrounded by WT heterozygous cells recognizable for their red ommatidia color due to the presence of p{W+] transgene on the sister WT chromosome. Notice the smaller and abnormally shaped mosaic eyes.

**Fig S5.**
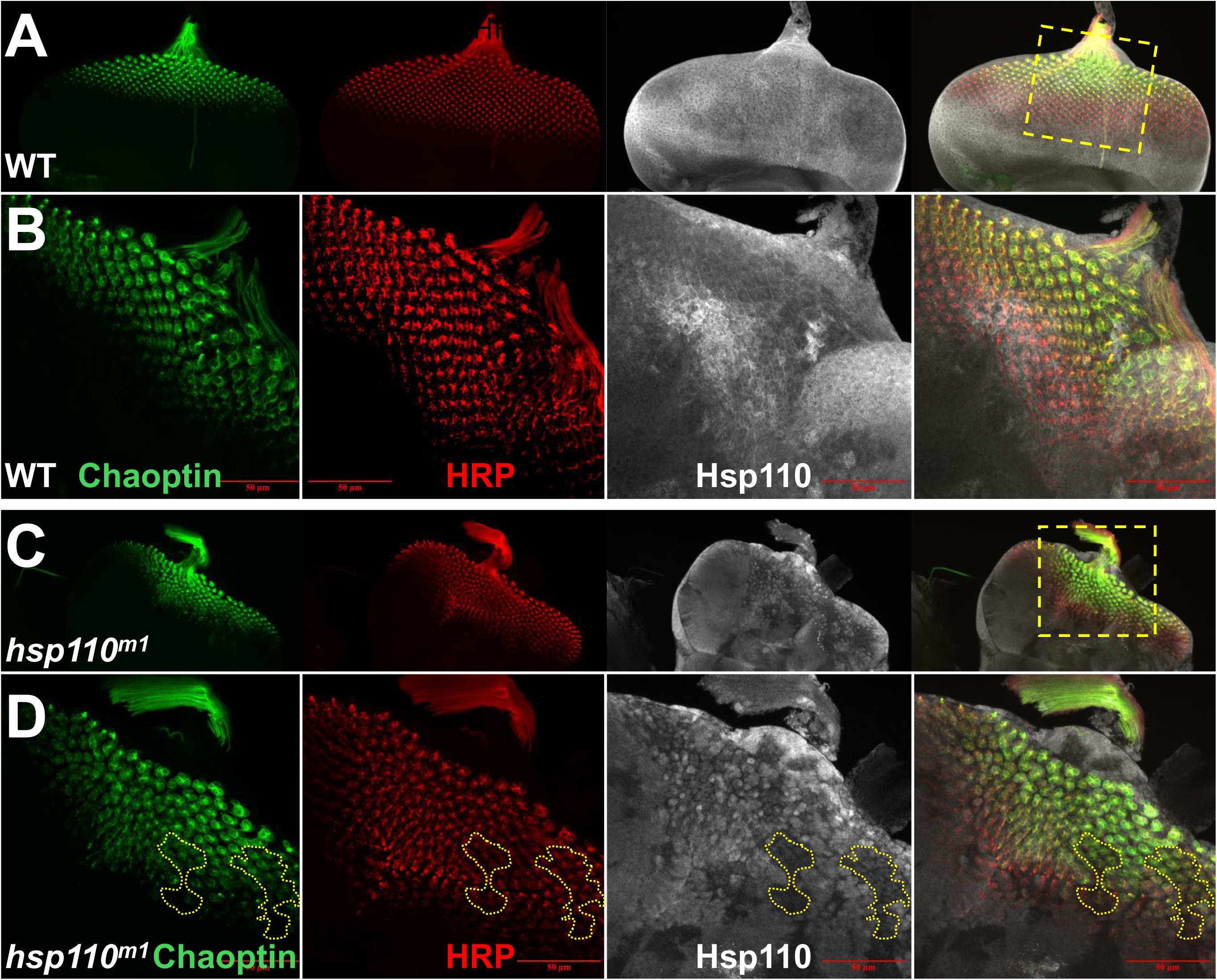
A non-cell autonomous role of Hsp110 for neuronal differentiation in developing eyes. Confocal images of eye imaginal discs from third-instar larvae of (A, B) WT control (genotype: *w^1118^*) or (C, D) carrying mosaic clones homozygous for *hsp110^m1^* mutant (genotype: *y, w, eyFLP2/+; M(3)RpS17^4^, P{w+}*, FRT80B/ *hsp110^m1^*, FRT80B), triple-labeled for Chaoptin (green), HRP (red) and Hsp110 (gray), shown in single channels or overlaying imaging, as annotated. (B, D) High-magnification views of regions highlighted with dashed yellow lines in (A) and (C), respectively. Yellow dashed lines in (D) highlight the clonal boundary of homozygous *hsp110^m1^* mutant cells, recognizable for their depleted Hsp110 levels as compared to surrounding heterozygous cells, that were also positive for neuronal markers Chaoptin and HRP, supporting that loss of endogenous Hsp110 does not block their differentiation into neuronal cell fate.

**Fig. S6.**
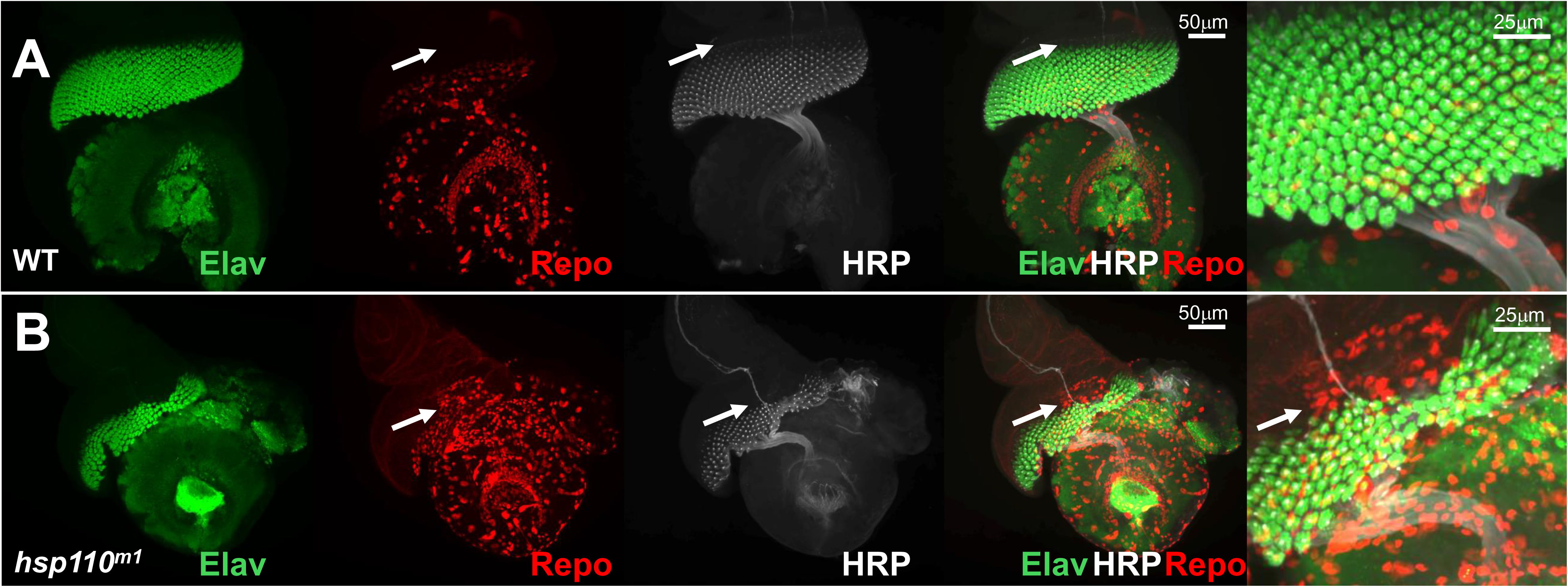
Hsp110 affects glial cell migration. Confocal images of brain and eye imaginal discs from third-instar larvae (A) of WT control (genotype: *w^1118^*) or (B) carrying mosaic clones for *hsp110^m1^* mutant (genotype: *y, w, eyFLP2/+; M(3)RpS17^4^, P{w+}*, FRT80B/ *hsp110^m1^*, FRT80B), triple-labeled for Elav (green), HRP (gray) and Repo (red), in single channels or overlaying imaging, as annotated. Glia cells normally migrate behind the front line of differentiating neurons in WT (arrows in A), but trespassed to the front area lacking neuronal differentiation (arrows in B) in an imaginal disc that contained large mutant clones.

## Materials and Methods

### Sequence analyses

Sequence alignment by Cluster V method using the LaserGene’s Megalign program, Hsp110 proteins from the following species were analyzed:

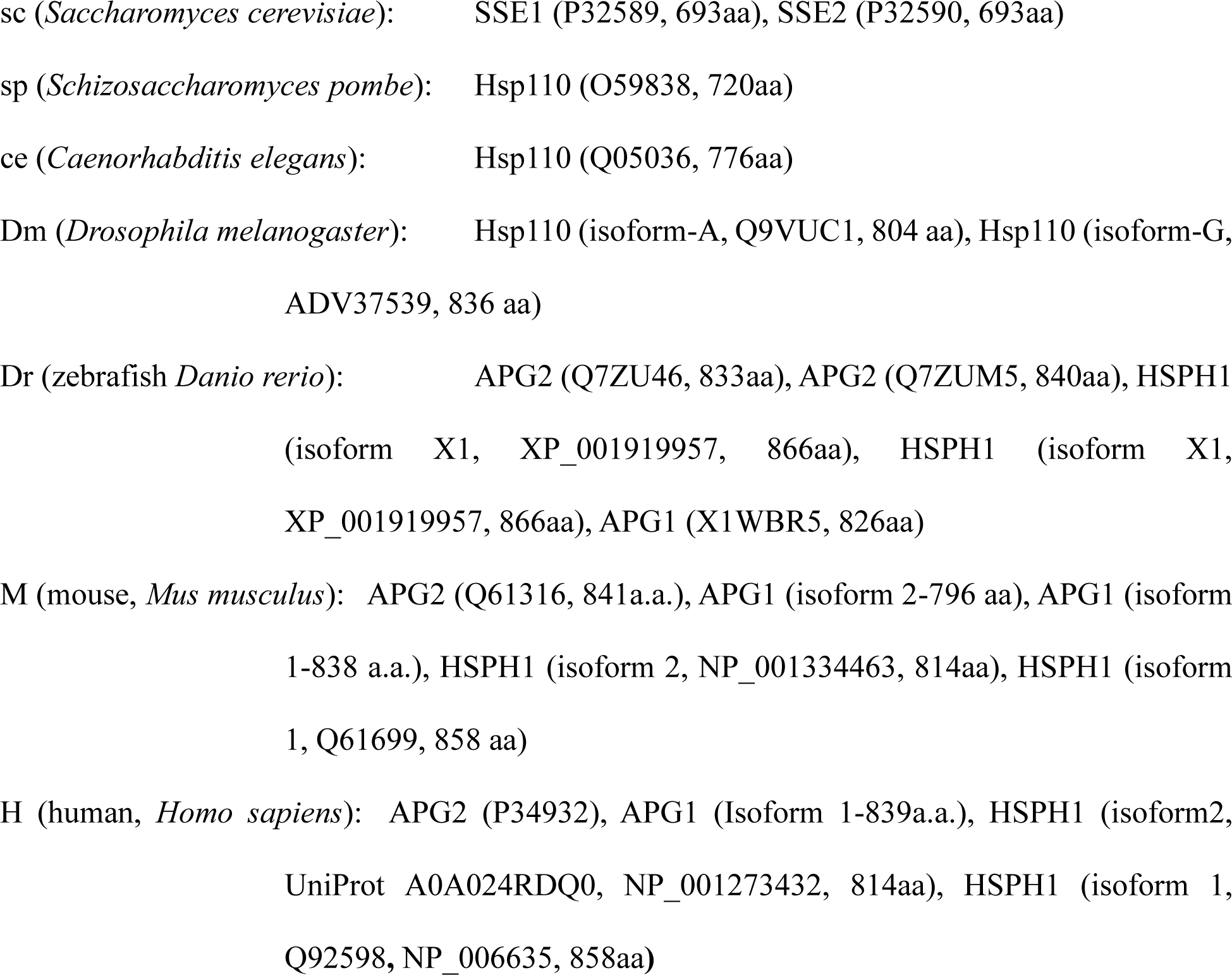

### *Drosophila* husbandry and genetics

Fly colonies were maintained at room temperature using standard husbandry procedures. Unless otherwise specified, all flies were reared at 25°C with 12 hrs dark and 12 hrs light cycle. Recombinant lines were created following standard fly genetics or as described (Xu and Rubin 1993). To remove the background mutations in *hsp110*^m1^ and *hsp110^FSVS1^* alleles, the two lines were crossed together and outcrossed for 5 generations.

### Drosophila stocks

Transgenic fly lines UAS-Hsp40 from Dr. Nancy Bonini (University of Pennsylvania). UAS-Hsp110 (isoform A) were described previously (Zhang et al. 2010) and verified by western blot analyses as well as immunofluorescent staining and confocal imaging. The following stocks were from Bloomington *Drosophila* Stock Center (BDSC): EY00671 (#15035); *hsp110^m1^* (*c00082 #*11485); *hsp110^m2^* (*l(3)s64906)*; Arm-GAL4 (#1560); daughterless(da)-Gal4 (#55850); Tubulin(Tub)-GAL4 (#5138); GMR-Gal4 (#9146); UAS-APG1 (#53729);UAS-luciferase RNAi (#31603);*w*\*; RpS17^4^ P{w(+mC)=arm-lacZ.V}70C, P{ry(+t7.2)=neoFRT}80B/TM6B, Tb^+^ (#6358); *y^d2^ w^1118^* P{ry(+t7.2)=ey-FLP.N}2 P{5xglBS-lacZ.38-1}TPN1; RpS17^4^ P{w(+m*) ry(+t7.2)=white-un1}70C P{ry(+t7.2)=neoFRT}80B/TM6B, P{y(+t7.7) ry(+t7.2)=Car20y}TPN1, Tb1 (# 5621); y1, w*; P{w(+mC)=tubP-GAL80}LL9 P{ry(+t7.2)=neoFRT}80B (#5191); MARCM analysis: P{ry(+t7.2)=hsFLP}1, P{w(+mC)=tubP-GAL4}1, P{w(+mC)=UAS-GFP.T:Myc.T:nls2}1, y1 w*; RpS174 P{w(+m*) ry(+t7.2)=white-un1}70C P{w(+mC)=tubP-GAL80}LL9 P{ry(+t7.2)=neoFRT}80B/TM6B, Tb1 (# 42732).*hsp110^FSVS1^* (KSC-115570) and *hsp110^FSVS2.^* (KSC-115275) were from Kyoto Stock Center.

### Bright field imaging of adult fly eyes

The heads of adult flies with appropriate genotypes were cut and orientated on glass slides with nail polish. Z-stack scanning of adult eyes were recorded under the 10X objective using Zeiss Axioimager Z1 microscope (usually 10-20 layers scanned) and reconstructed into 3D projection using CZFocus software.

### Dissection and staining of adult eye retina

Eye retina dissection was performed as described (Nie, Mahato, and Zelhof 2015). Briefly, female flies of a specific genotype were beheaded. Then their heads were torn open from their proboscis to let 4% paraformaldehyde go in. After those heads were fixed for 1 hour at room temperature, they were washed briefly with PBS twice. Then their retinas were carefully removed from the corneal lens layer through fine forceps and a tungsten hook. The dissected retinas were washed with PBS three times, and stained with Alexa Flour 488 Phalloidin (1:1000, Molecular Probes, Cat#: A12379) overnight at 4 °C. The next day, the samples were washed with PBS, mounted, and imaged using a confocal microscope (Leica DM6000-B).

### Embryo immunofluorescent staining

About 300 flies were placed on apple juice plates to lay eggs overnight. Embryos were collected and then treated with 50 percent bleach diluted in water for 2 minutes. They were rinsed with water for a few minutes and then fixed in a 4% PFA/heptane (1:1) mixture for 30 minutes. After the lower phase of the fixative was removed and equal parts of methanol were added, the embryos were shaken for 30 seconds. Then the upper liquid (also for the embryos that did not sink) was removed and the embryos were rinsed with methanol two more times. Before being stained by primary antibody, the embryos were rehydrated sequentially by different methanol/PBT (2:1, 1:1, 1:2) mixtures for 5 min each. Then embryos were washed by PBT for 15 mins before blocked with 5% normal goat serum.

### *Drosophila* whole-mount immunofluorescent staining

Fly ovaries, larval neuromuscular junctions (NMJs), brains, and imaginal discs were dissected in ice-cold 1XPBS and fixed with 4% paraformaldehyde (PFA) for 30 minutes to 1 h at room temperature. After fixation, tissues were rinsed with 1XPBS, washed with 1XPBT (1XPBS with 0.3% Triton X-100), and blocked with 5% normal goat or donkey serum diluted in 1XPBT (i.e., blocking solution) for 1 h at room temperature. Tissues were then incubated overnight at 4 °C with primary antibodies diluted in the blocking solution, washed with 1XPBT, and incubated with secondary antibodies diluted in 1XPBT for 2h at room temperature or overnight at 4°C. After secondary incubation, the samples were washed in 1XPBT and counterstained with DAPI (1:5000 dilution), rinsed with 1XPBS. Samples were then mounted with antifade mounting medium (H-1900, Vector Laboratories), and imaged on a Nikon AX-R confocal microscope.

### Western blotting

#### Protein extraction

Whole-animal homogenates were prepared by collecting adults in 1.5 mL Eppendorf tubes and gently homogenized with ice-cold RIPA lysis buffer (0.5% NP-40, 0.25% sodium deoxycholate, 0.1% SDS, 150 mM KCl, 1 mM EDTA, 20 mM Tris (pH 7.5) containing protease inhibitors (GenDepot). Homogenates were centrifuged at 12,000 rpm for 10 minutes at 4°C. Supernatants were collected, boiled with SDS for 5 minutes, and then immediately stored in −80°C or separated on 8% SDS-PAGE gels and transferred onto nitrocellulose membranes (Millipore) overnight at 4°C. Membranes were rinsed with 1X Tris-buffer saline containing 0.1% Tween-20 (1XTBST), blocked with 1.5% casein for 1 h at room temperature, and then incubated with primary antibodies overnight at 4 °C. The following day, the membranes were washed with 1XTBST and incubated with IRDye® 680/800 secondaries for 1 h at room temperature, washed, and resolved using an LICOR Odyssey Infrared Imaging System.

### Western Quantification

Band intensities were quantified and normalized to the corresponding Actin or Tubulin loading controls for each lane using ImageJ/Fiji software (NIH). The signal-to-loading control ratio for the WT lane was set equal to 1, and the relative fold change in expression was determined by diving the signal-to-loading control ratio for each sample by the WT ratio.

### Anti-Drosophila Hsp110 antibodies

cDNA for full-length *Drosophila* Hsp110 isoform A (LD32979, Drosophila Genomics Resource Center) with an in-frame 6xHis-tag in pET16b vector was purified from E. coli and used to immunize three rats (Pocono Rabbit Farm & Laboratory, Inc., Canadensis, PA). Crude sera were diluted to a final working concentration of 1:500-1:1,000 for Western blot and 1:200 for immunofluorescent staining.

### Antibodies

#### Primaries used

Chicken anti-GFP (Aves, 1:5000 for western blot and 1:1000 for staining); Rat-anti-Eav (7E8A10), mouse anti-Elav (9F8A9), Repo (8D12) and Chaoptin (24B10) were from Developmental Studies Hybridoma Bank (DSHB) and used at 1:50 for immunofluorescent staining. Mouse anti-Actin (1:10000, MAB1501, Chemicon, and AB6276, Abcam).

#### Secondaries used

Alexa Fluor 488 AffiniPure Donkey anti-Chicken IgY (1:1000, 703-545-155, Jackson ImmunoResearch Inc), Alexa Fluor® 488 AffiniPure™ Donkey Anti-Mouse IgG (1:1000, 715-545-150, Jackson ImmunoResearch Inc), Rhodamine Red AffiniPure Donkey anti-Mouse IgG (1:1000, 715-295-151, Jackson ImmunoResearch Inc), DyLight 549 Goat anti-Mouse IgG (610-142-002, Rockland Immunochemicals Inc), Alexa Fluor 594 AffiniPure Donkey Anti-Mouse IgG (715-585-151, Jackson ImmunoResearch Inc), Alexa Fluor® 647 AffiniPure Donkey Anti-Rat IgG (712-605-153, Jackson ImmunoResearch Inc), Rhodamine-X conjugated Goat-anti-HRP (Jackson Laboratories), TRITC-Phalloidin (1:200, P1951, Sigma), Alexa Fluor 647-Phalloidin (1:200, A-30107). Alexa Fluor 680 Goat anti-Rabbit IgG (1:5000, A21076, Invitrogen), IRDye® 680RD Goat anti-Mouse IgG (1:5000, 926-68070, LI-COR), IRDye® 800CW Donkey anti-Mouse IgG (1:5000, 926-32212, LI-COR).

### Data Collection and Statistical Analysis

More than 10 samples were analyzed for each of the studied genotypes. All datasets were first evaluated for normality using the Shapiro-Wilk test. Pairwise two-sample Kolmogorov-Smirnov (KS) tests with Benjamini-Hochberg false discovery rate (FDR) correction were used to compare differences in data distributions. Significance was defined as: P > 0.05 as non-significant (ns), *P < 0.05, **P < 0.01, ***P < 0.001, and ****P < 0.0001, with the defined N described in the corresponding figure legends.

## Acknowledgments

We thank the Bloomington *Drosophila* Stock Center (BDSC) for providing *Drosophila* lines and the Developmental Studies Hybridoma Bank (DSHB) for supplying *Drosophila* antibodies, Dr. Zhengmei Mao at the IMM Microscopy Core of UTHealth Houston at the Center for Advanced Microscopy (CAM) of UTHealth Houston for support with confocal microscopy and structure-illumination microscopy assistance, respectively, and the Bovay Foundation for funding support.

